# Next-generation insect digitization: combining phenomics and genomics by subsequent synchrotron X-ray imaging and DNA sequencing

**DOI:** 10.64898/2026.08.20.745929

**Authors:** Cristina Lupașcu-Vasilița, Alexander Riedel, Daniela Mera-Rodríguez, Angelica Cecilia, Tomáš Faragó, Elias Hamann, Jenny Hein, Annette Herz, Jakob Martin, Janes Odar, Pauline Pfeiffer, Chandan Sarkar, Rebecca Spiecker, Clément Tavakoli, Marcus Zuber, Christian Rabeling, Tilo Baumbach, Lars Krogmann, Thomas van de Kamp

**Affiliations:** Institute for Photon Science and Synchrotron Radiation (IPS), Karlsruhe Institute of Technology (KIT), Hermann-von-Helmholtz-Platz 1, 76344 Eggenstein-Leopoldshafen, Germany; Department of Entomology, State Museum of Natural History Stuttgart, Rosenstein 1, 70329 Stuttgart, Germany; Institute of Biology, Biological Systematics (190w), University of Hohenheim, Garbenstr. 30, 70599 Stuttgart, Germany; State Museum of Natural History Karlsruhe, Erbprinzenstr. 13, 76131 Karlsruhe, Germany; Institute of Biology, Department of Integrative Taxonomy and Biodiversity of Insects, University of Hohenheim, Garbenstr. 30, 70599 Stuttgart, Germany; Center for Biodiversity and Integrative Taxonomy Research, University of Hohenheim & State Museum of Natural History Stuttgart, Stuttgart, Germany; Institute for Biological Control, Julius Kühn Institute (JKI)-Federal Research Centre for Cultivated Plants, Schwabenheimer Str. 101, 69221, Dossenheim, Germany; Laboratory for Applications of Synchrotron Radiation (LAS), Karlsruhe Institute of Technology (KIT), Kaiserstr. 12, 76131 Karlsruhe, Germany

## Abstract

Recent technological advances allow for the large-scale acquisition of genetic and morphological data: high-throughput sequencing has transformed the field of genomics while synchrotron X-ray microtomography enables rapid, noninvasive 3D imaging. However, integrating these approaches for the same specimens is challenging because X-rays can fragment DNA, and DNA extraction damages internal morphology, particularly relevant for small bodied organisms, such as insects. We systematically tested multiple extraction protocols and irradiation conditions across three model insect species. We irradiated more than 1,000 specimens under varying conditions and tested DNA quality through DNA barcoding and UCE sequencing. Our results demonstrate that high-quality DNA and high-resolution tomograms can be obtained from the same individuals, provided that the parameters are carefully optimized and rapid SR-μCT scanning precedes DNA extraction. In this respect, our findings establish practical guidelines for combining genomics and phenomics, paving the way for comprehensive integrative digitization of biodiversity.

## Introduction

The digital transformation of our society greatly enhances the possibilities to study biodiversity at an unprecedented scale, as extensive genetic and morphological data can be generated, analyzed and made available online worldwide^1^. Insects are by far the most diverse group of animals on Earth, representing a significant share of global biodiversity. They perform key ecosystem functions^2^, are critical for agriculture^3^, serve as models for biomimetic design^4^, are indicators of biodiversity decline^5^ and climate change^6^ and have become an integral part of genetic research and developmental biology^7^. Their ecological importance, combined with their immense species diversity and comparatively small body size, positions insects at the forefront of biodiversity research and development of advanced technologies far large-scale digitization approaches.

Genetic sequencing revolutionized biodiversity discovery and biomonitoring^8^, research on population structure and speciation^9^, as well as phylogenetics^10,11^. Genetic barcoding has been widely used for species identification as well as the reconstruction of shallow phylogenetic relationships^12^ and has recently resurfaced as a great tool for rapid, high-throughput and large-scale biodiversity discovery^13^. In recent years, insect genomics has been greatly enhanced by the advent of next generation sequencing technologies. Long read sequencing improved the quality of genome assemblies and the resolution of complex and repetitive regions^14,15^, providing insights into the evolution and rapid adaptation of insects in the context of climate change and biodiversity loss^16^. Additionally, reduced-representation genome sequencing (RRGS) approaches are now able to provide large-scale multigene/multilocus data sets^17^ for phylogenetics, population genetic studies or conservation genetics^18^. The state-of-the-art RRGS method is capturing and sequencing ultraconserved elements (UCEs), highly conserved gene regions that are scattered throughout the genomes of most organisms and flanked by regions of greater sequence variability^19^. UCE sequencing enables the reconstruction of evolutionary history across different phylogenetic levels^20,21,22^. The outlined technological advancements paved the way for *high-throughput genomics*^23^, allowing the large-scale acquisition of genetic data from entire insect collections.

In parallel, digitization of insect morphology has become a major effort worldwide. Although the digitization of large collections is often restricted to simple 2D (stack-) photography^24^, 3D imaging via photogrammetry^25^ can be applied to create detailed volumetric models that preserve surface texture. In order to digitize specimens as 3D volumes that capture both external and internal morphological information in high-resolution, X-ray microtomography (µCT), is becoming increasingly popular. The technique allows tackling a wide range of scientific questions, ranging from physiology^26,27^, evolution^28,29^, paleontology^30,31^, ecology^32^ over comparative morphology^33^ to functional morphology^34^ and biomimetics^35^. Compared to laboratory X-ray sources, synchrotron light sources offer much higher flux density and coherence^36^. Phase contrast options enable enhancing soft tissue contrast in ethanol-preserved specimens, often making staining unnecessary^37,38^. Acquisition times per sample can be reduced to below one minute^39^, allowing synchrotron radiation microtomography (SR-µCT) to be applied to series of hundreds and even thousands of specimens^28,32,39^. It is likely to become the gold standard for *high-throughput phenomics*, enabling the digitization of morphological diversity across entire collections of insects and other small organisms^40^.

Both genome sequencing and synchrotron X-ray imaging have independently driven recent substantial advances in entomology, contributing to a “digital revolution” in the field. However, an immense opportunity lies in the systematic integration of both approaches at large scale by combining *high-throughput genomics* and *high-throughput phenomics*. Huge collections of paired datasets of genetic and morphological information will open up the possibility of mapping morphological characters onto molecular phylogenies to identify evolutionary key events and even facilitate systematic genotype-phenotype correlations. The ongoing increase in automation of both imaging and data analysis^40,41^ offers enormous potential for comprehensive and multidimensional analyses of insect biology, which will likely accelerate with the use of AI, an advance that could be transformative for entomology as a whole.

Although sequencing and imaging different conspecific individuals can be an option, the ideal solution for insect digitization, arguably lies in acquiring paired morphological 3D and genome data from the same individual. Such an approach accounts for intraspecific variation, precise phenotype-genotype correlations and ultimately the possibility to study evolution of genes and their functions. It eliminates confusion with similar and cryptic species, provides digital vouchers for long-term validation and cross-referencing, and therefore is scientifically rigorous and future-proof.

However, such an integrative approach presents considerable challenges. On the one hand, X-rays are ionizing and known to damage DNA^42,43,44^. The effects include mutations, breaks of DNA strands, base modification and structural changes such as gene order rearrangements. Oxidative damage can cause strand fragmentation and X-ray-induced free radicals may lead to strand breaks, resulting in smaller fragments^45^.

On the other hand, DNA extraction typically destroys internal anatomy, especially soft tissues. This is particularly notable in proteinase K-based extraction methods, where the enzyme breaks down muscle tissue. Although it is common practice in insect research to extract a small tissue sample (e.g. a leg) for DNA analysis, this is often extremely difficult for small insects and typically yields insufficient genetic material. In addition, even a seemingly minimal intervention, such as removing a leg or part thereof, causes irreversible damage to the specimen’s anatomy. Moreover, this approach is unsuitable for large-scale studies due to the time-intensive nature of the dissection and the increased risk of cross-contamination. Other options of widely used DNA extraction methods include DNA leaching with an alkaline solution, for example the HotSHOT buffer. Since there is no digestive enzyme involved, such “gentle” DNA leaching methods are generally considered to be non-destructive to the soft tissues^46,47^.

The prevailing assumption in the community was that DNA sequencing after SR-µCT is not feasible. Therefore, “gentle” DNA extraction before SR-µCT seemed to be the more viable option. However, we found that even “gentle” methods caused internal morphological damage. This observation raised two central questions: First, are there DNA-extraction methods that minimize internal morphological damage to a degree that still permits meaningful morphological analysis? And second, if not, can efficient X-ray imaging sufficiently limit DNA degradation to allow subsequent DNA sequencing?

Using DNA barcoding and UCE sequencing in concert with high-throughput SR-µCT, we analyzed whether it is possible to apply both techniques sequentially without compromising data quality, and if so, which order is preferable. Two conditions were defined as successful integration: tomographic scans must be of sufficient quality for morphological analysis, and the quality of the DNA must be adequate for DNA sequencing.

For our study, we selected insect species representing three of the largest insect orders: Coleoptera, Diptera, and Hymenoptera. The chosen species span a range of body sizes and sclerotization: the compact and heavily sclerotized grain weevil (*Sitophilus granarius*), the soft-bodied spotted wing drosophila (*Drosophila suzukii*), and the slender and sclerotized wasp *Leptopilina japonica* (Fig. 1). *Sitophilus granarius* and *D. suzukii* are major agricultural pests, while *L. japonica* is an important parasitoid of *D. suzukii*.

**Fig. 1.**
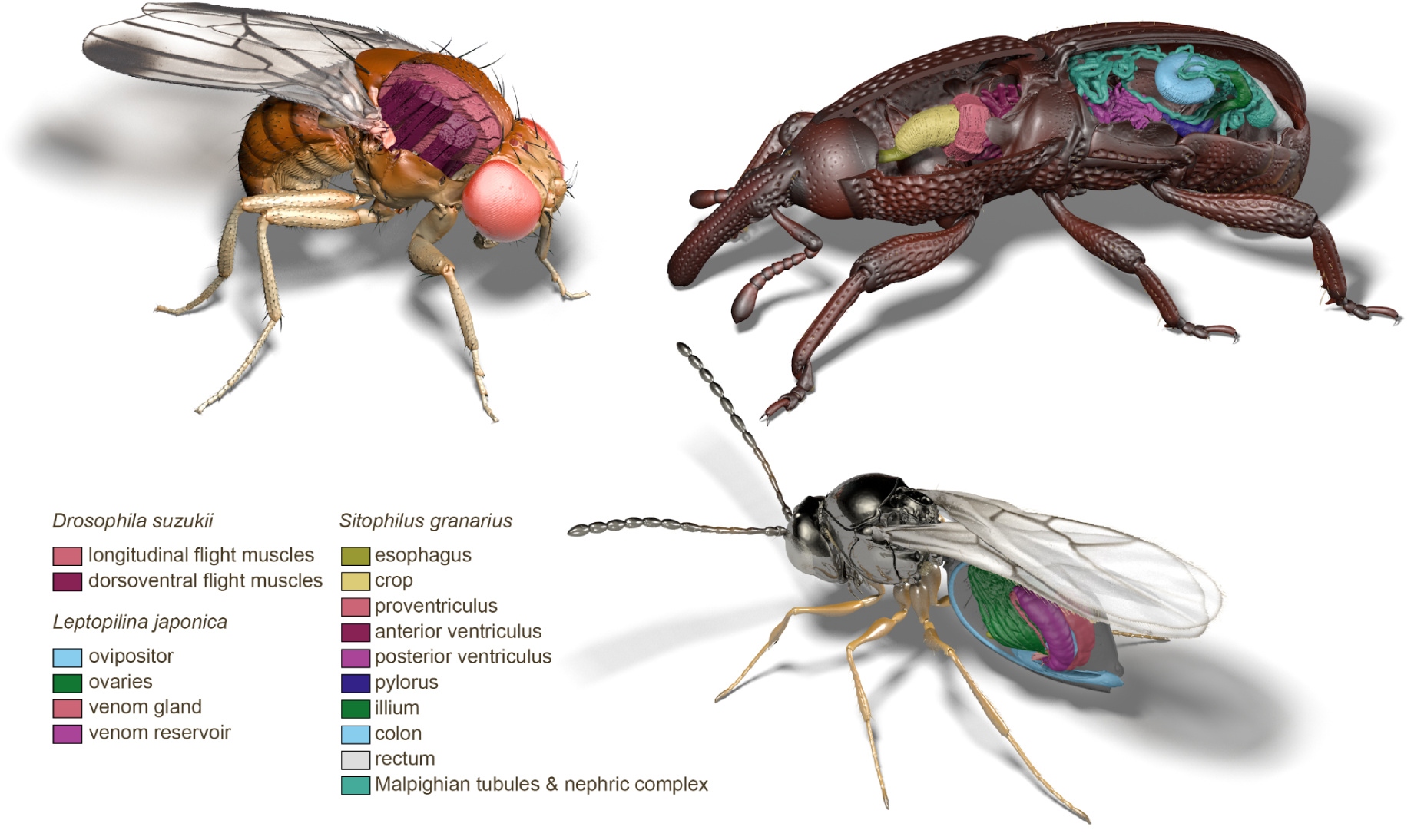
Three-dimensional renderings of virtually sectioned insects based on synchrotron X-ray microtomography. Surface models composed of individual body parts segmented from 3D data acquired in this study, showing the three examined species digitally cut with partially exposed internal structures: *Drosophila suzukii* (left) with longitudinal and dorsoventral flight muscles, *Sitophilus granarius* (top right) with the alimentary canal, and *Leptopilina japonica* (bottom right) with the female reproductive system and venom apparatus. False colors indicate internal anatomical structures, while the external body colors were added artificially for a more natural appearance.

Our experimental design comprised two complementary experiments (Fig. 2). First, we investigated the effects of five different DNA extraction protocols on internal insect morphology. To directly quantify extraction-induced damage, the same individuals were scanned tomographically before and after DNA extraction. In a second experiment, we examined X-ray-induced DNA degradation as a function of irradiation time and beam parameters (polychromatic vs. monochromatic) and the compatibility of DNA sequencing approaches with irradiated material, addressing whether tomographic datasets of sufficient quality can be acquired without excessive DNA degradation.

**Fig. 2.**
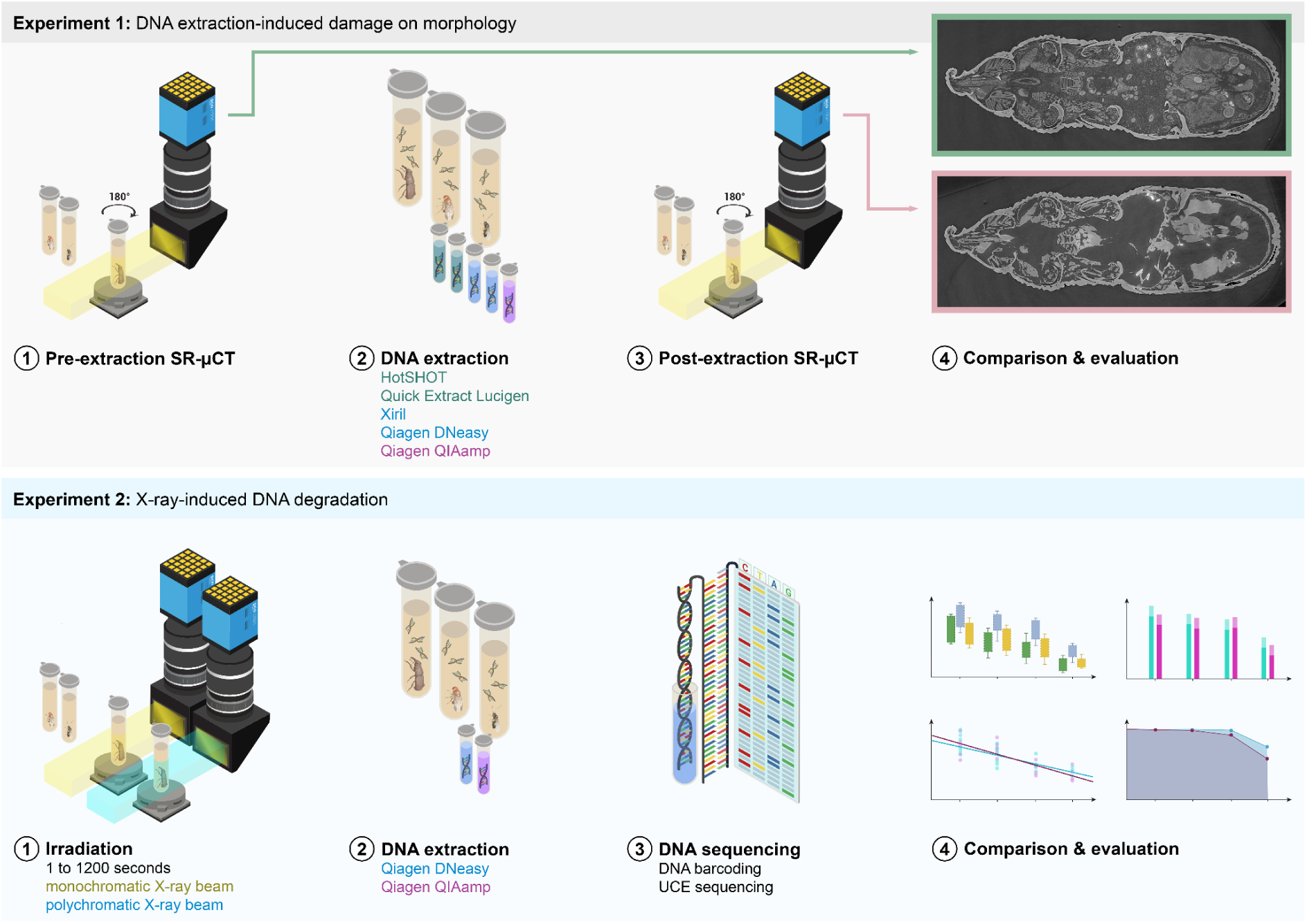
Overview of the two complimentary experiments. In experiment 1, the same individual insects were scanned before and after DNA extraction. Afterwards, the morphological damage caused by five different DNA extraction protocols was evaluated by comparing pre- and post-extraction tomograms. In experiment 2, insects were irradiated between 2 and 1,200 seconds using a monochromatic or polychromatic X-ray beam. Afterwards, DNA was extracted from the irradiated specimens using Qiagen DNeasy or Qiagen QIAamp DNA extraction protocols, followed by DNA sequencing (barcoding and UCEs) and analysis of DNA concentration and fragmentation.

Our results provide practical guidelines for a true integration of genomic and phenomic insect data, particularly for large-scale, multimodal digitization initiatives aiming to combine *high-throughput genomics* and *high-throughput phenomics*.

## Results

### Morphology before DNA extraction

Prior to DNA extraction, the tomograms generally showed a well-preserved morphology and the various soft tissues such as muscles, nervous system, digestive tract and genitalia were well defined. In *D. suzukii* (Figs. 3C, 4I, 6A,C,E,G,I and S2A,C,E,G,I) ethanol-fixation resulted in some shrinkage of the muscles and many individuals showed a collapsed cuticle, especially of the compound eyes. In *S. granarius* (Figs. 3A, 4A,E, 5A,C,E,G,I, S1A,C,E,G,I and S4) and *L. japonica* (Figs. 3E, 4C,G, 7A,C,E,G,I and S3A,C,E,G,I), soft tissues remained intact and no pronounced signs of shrinkage or deformation were detected. However, in some individuals of *S. granarius* a contraction of the dorsal abdominal region was noted.

**Fig. 3.**
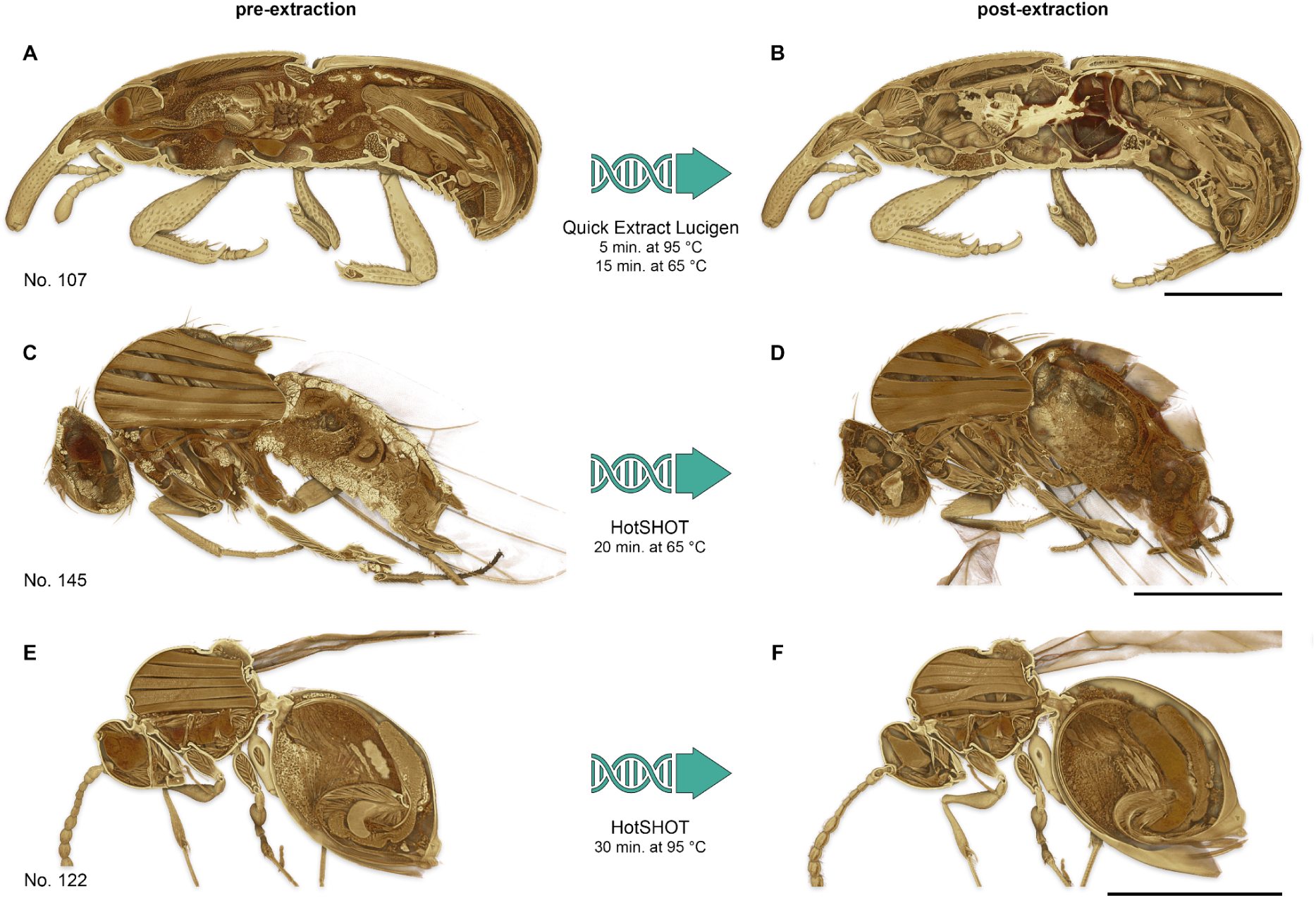
Sectioned volume renderings of insect specimens before and after “gentle” DNA extraction. Examples show the same individuals scanned pre- and post-extraction: *Sitophilus granarius* (A,B), *Drosophila suzukii* (C,D), and *Leptopilina japonica* (E,F). Before extraction, the internal anatomy is well preserved, whereas post-extraction scans reveal varying degrees of tissue shrinkage and deformation. Scale bars = 1 mm.

**Fig. 4.**
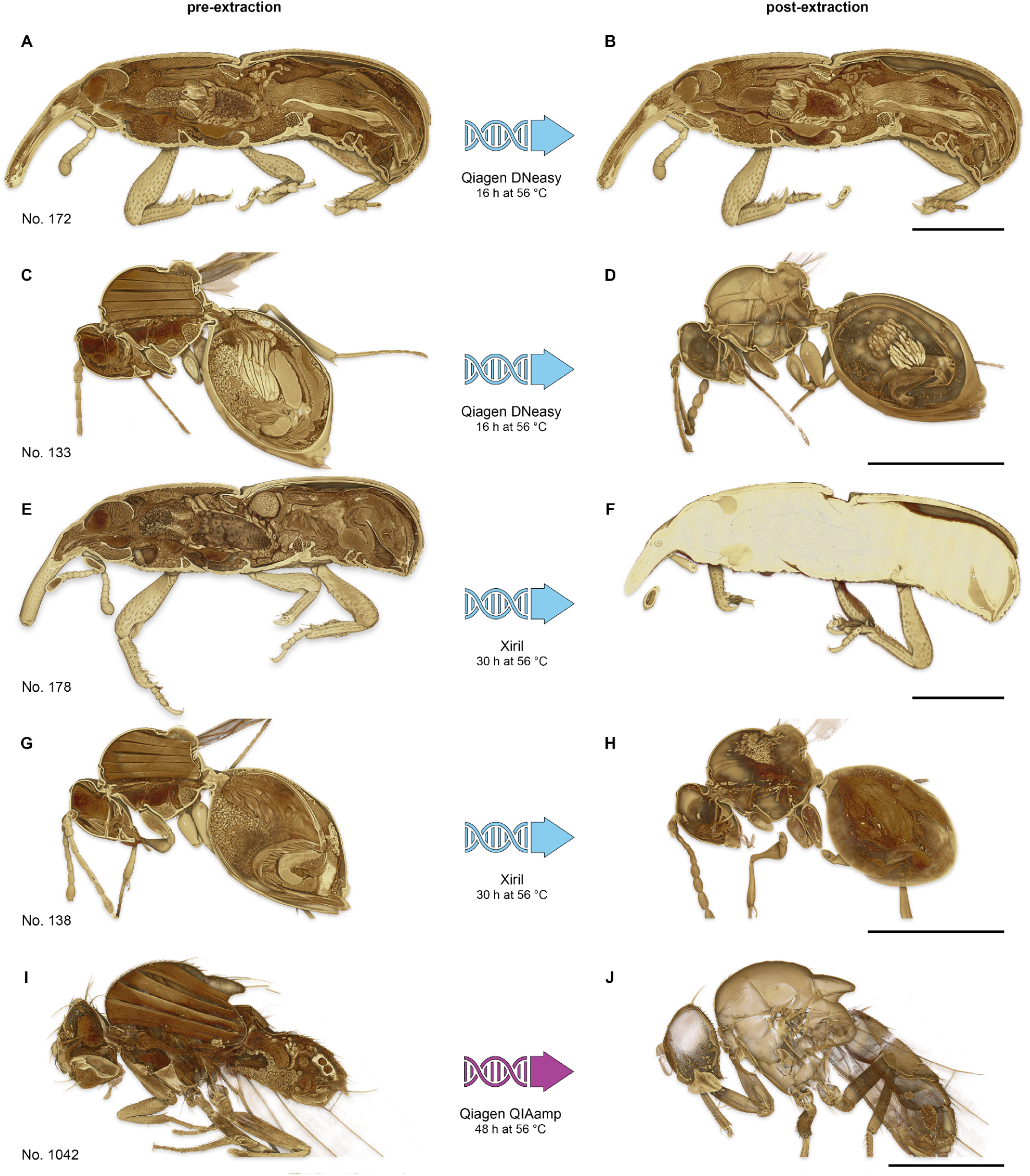
Sectioned volume renderings of insect specimens before and after proteinase K-based DNA extraction. Examples show the same individuals scanned pre- and post-extraction using different protocols highlighting species-specific effects. Using *Qiagen DNeasy*, *Sitophilus granarius* (A,B) showed slightly increased X-ray absorption but largely preserved anatomy; the soft tissues of *D. suzukii* and *L. japonica* (C,D) were largely digested, the latter retained only the ovaries. *Xiril* extraction caused strong X-ray absorptivity in *S. granarius* (E,F) with most internal organs remaining largely intact, but led to soft tissue digestion in *D. suzukii* and *L. japonica* (G,H). *Qiagen QIAamp* resulted in complete digestion of internal anatomy in *D. suzukii* (I,J) and *L. japonica*, but had little effect on *S. granarius*. Scale bars = 1 mm.

**Fig. 5.**
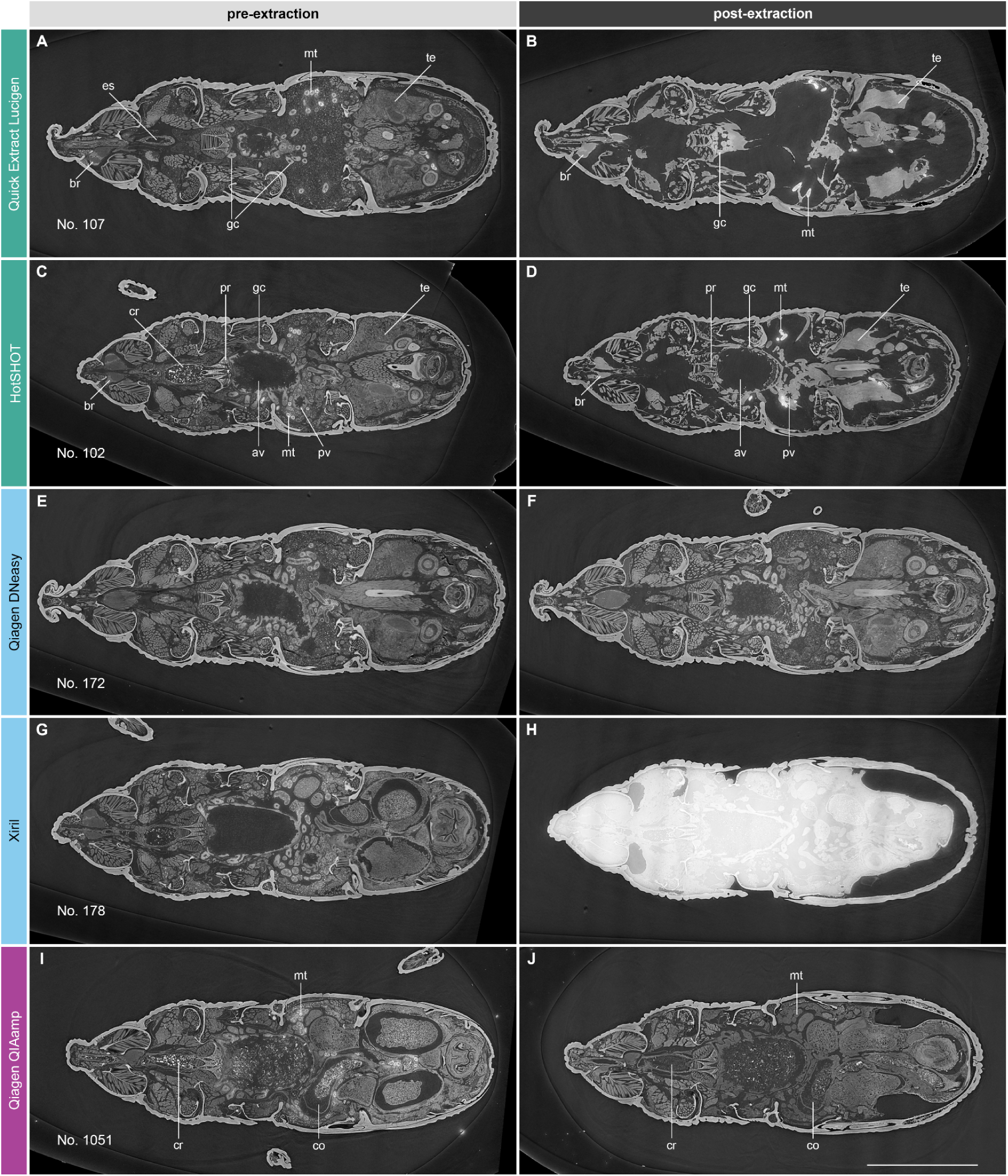
*Sitophilus granarius* before and after DNA extraction. Representative horizontal tomographic slices highlight the effects of different DNA extraction methods on internal anatomy. av = anterior ventriculus; br = brain; cr = crop; co = gut content; es = esophagus; gc = gastric caeca; mt = Malpighian tubule; pr = proventriculus; pv = posterior ventriculus; te = testis. Scale bar = 1 mm.

**Fig. 6.**
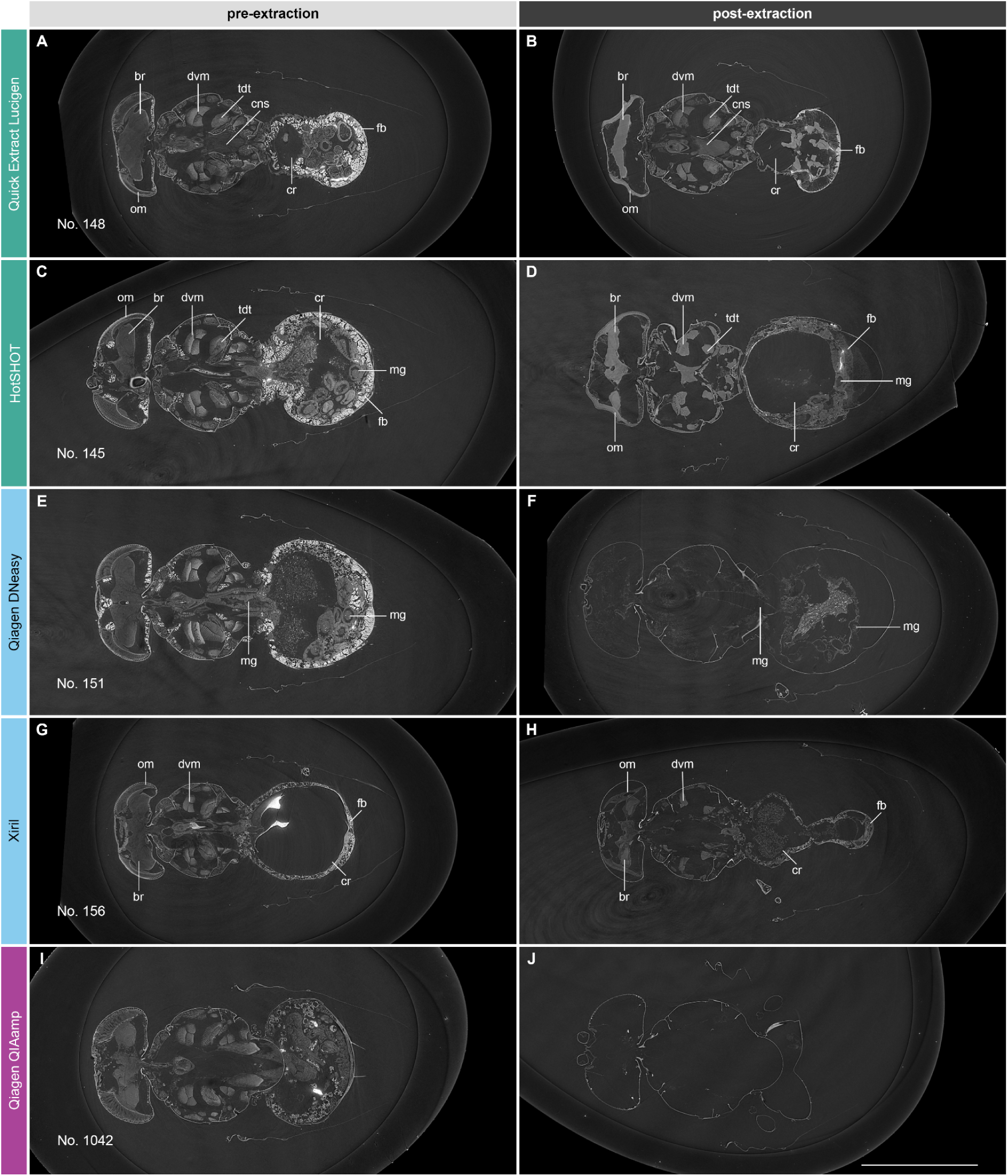
*Drosophila suzukii* before and after DNA extraction. Representative horizontal tomographic slices highlight the effects of different DNA extraction methods on internal anatomy. br = brain; cns = central nervous system; cr = crop; dvm = dorsoventral flight muscle; fb = fat body; mg = midgut; om = ommatidia; tdt = tergal depressor of the trochanter. Scale bar = 1 mm.

**Fig. 7.**
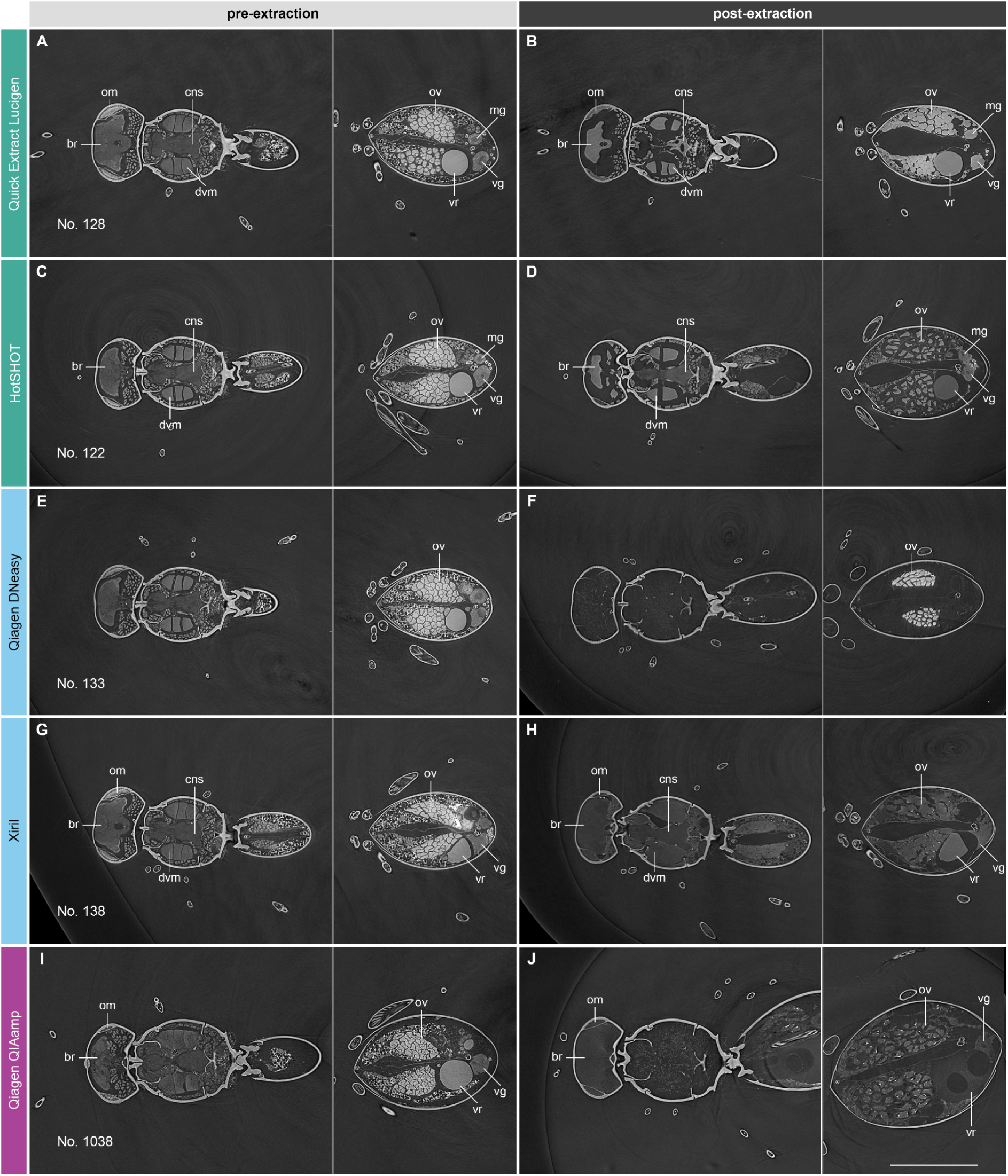
*Leptopilina japonica* before and after DNA extraction. Representative horizontal tomographic slices highlight the effects of different DNA extraction methods on internal anatomy. br = brain; dvm = dorsoventral flight muscle; cns = central nervous system; mg = midgut; om = ommatidia; ov = ovary; vg = venom gland; vr = venom reservoir. Scale bar = 0.5 mm.

### Morphology after DNA extraction

Although individuals of a given species exhibited similar anatomical effects when treated with a particular DNA extraction protocol, we observed striking interspecific differences for the “proteinase K-based protocols”, particularly between *S. granarius* and the other two study species.

The “gentle” <u>Quick Extract Lucigen</u> (QE) treatment heavily altered the internal anatomy of all three species. Muscles and soft tissues generally shrank considerably. In *S. granarius* (Figs. 3B, 5B, S1B and Movie S1), several organs could be identified after treatment, including the brain, parts of the alimentary canal, and the testes. However, major parts of the internal anatomy were ruptured and destroyed. Malpighian tubules showed stronger X-ray absorptivity. Similar effects were observed in the soft tissues of *D. suzukii* (Figs. 6B, S2B and Movie S2). The cuticle sometimes collapsed, especially of the eyes. Organs that could be identified after treatment included the brain, central nervous system, crop, and fat body. The fat body appeared smaller and showed less X-ray absorptivity. *Leptopilina japonica* (Figs. 7B, S3B and Movie S3) exhibited shrinkage similar to that observed in the other two species. Among the organs that could be identified after treatment were the brain, central nervous system, and most parts of the genitals.

The effects of the second “gentle” approach, <u>HotSHOT</u> (HT), were comparable with respect to soft tissue shrinkage and the ability to identify certain organs. In *S. granarius* (Figs. 5D, S1D and Movie S1), the increased X-ray absorptivity of the Malpighian tubules was noteworthy again. In *D. suzukii* (Figs. 3D, 6D, S2D and Movie S2), the cuticle again showed collapse, and the crop was sometimes bloated. In *L. japonica* (Figs. 3F, 7D, S3D and Movie S3), the lower X-ray absorptivity of the genitals was notable. In some individuals, the abdomen was bloated, and the follicles of the ovaries were dispersed.

The proteinase K-based <u>Qiagen DNeasy</u> (Qi) protocol had almost no effect on the anatomy of *S. granarius* (Figs. 4B, 5F, S1F and Movie S1), though slightly increased X-ray absorptivity was observed throughout the specimens. In contrast, the soft tissues of *D. suzukii* (Figs. 6F, S2F and Movie S2) were largely digested, resulting in an empty shell. Only very faint remains of some organs, such as the gut and crop, could sometimes be observed. As with the latter species, the soft tissues of *L. japonica* (Figs. 4D, 7F, S3F and Movie S3) were digested. Notably, the ovaries shrank but remained clearly visible after treatment.

<u>Xiril</u> (Xi), the second proteinase K-based protocol tested in this study, had a striking effect on *S. granarius* (Figs. 4F, 5H, S1H and Movie S1), as all of the weevils treated with this method exhibited a much higher X-ray absorptivity. Otherwise, we only observed some collapse of the abdomen. Most of the organs remained largely intact. In *D. suzukii* (Figs. 6H, S2H and Movie S2), the protocol resulted in severe damage and collapse of soft tissues; yet, some organs, such as the brain, ommatidia, crop, and fat body, could still be identified. *Leptopilina japonica* (Figs. 4H, 7H, S3H and Movie S3) showed variable preservation, with some individuals retaining partially intact structures. The digestive effect was clearly visible, though, and while some organs were identifiable, there was a visible loss of contrast between different structures. Similar to the HotSHOT treatment, the genitals exhibited decreased X-ray absorptivity.

The third proteinase K-based extraction method tested was the <u>Qiagen QIAamp</u>, adjusted to target DNA recovery from museum specimens (MU). The protocol had only minor effects on *S. granarius* (Figs. 5J, S1J and Movie S1). We merely observed some shrinkage of the abdomen. Notably, there was decreased X-ray absorptivity of certain organs, such as the Malpighian tubules, as well as the crop and gut contents. For *D. suzukii* (Figs. 4J, 6J, S2J and Movie S2), the protocol yielded completely digested soft tissues, leaving behind empty, partially collapsed shells. In *L. japonica* (Figs. 7J, S3J and Movie S3), preservation ranged from damage and collapse of internal structures to complete digestion. In individuals with lesser effects, though heavily deformed and partially digested, some organs, such as the brain, ommatidia, and parts of the genitals, could still be identified. Sometimes, the abdomen was bloated.

### Deposited radiation dose

Dose simulations for SE2 (Methods) revealed that all three species experienced comparable dose rates under monochromatic irradiation, ranging from 189 ± 21 Gy/s in *Drosophila suzukii* and *Leptopilina japonica* to 191 ± 21 Gy/s in *Sitophilus granarius* (Table S1). In contrast, exposure to the filtered white beam resulted in an approximately 14-fold increase in dose rate, reaching 2,742–2,770 Gy/s across species. Simulated cumulative doses increased accordingly with exposure time, ranging from 379 ± 46 Gy after 2 s to 227.6 ± 25.4 kGy after 1200 s in monochromatic mode, and from 5.5 ± 0.7 kGy to 3,303.8 ± 369.4 kGy for the filtered white beam (Table S2).

### DNA concentration in tomographed samples

DNA concentration varied substantially among taxa and extraction methods in the SE1 samples. Under the HT protocol, *S. granarius* showed the highest mean concentrations and the greatest variability, whereas *L. japonica* had the lowest concentrations. In contrast, with QE extraction, *S. granarius* exhibited the lowest mean concentration and smallest standard deviation, while *D. suzukii* yielded the highest concentrations; a similar pattern was observed for Qi extraction. Under both Xi and MU protocols, *D. suzukii* again produced the highest mean concentrations with relatively high variability, whereas *S. granarius* consistently showed the lowest overall concentrations. Across methods, *L. japonica* displayed comparatively low variability (Table S3).

### DNA concentration in irradiated samples

DNA concentration was primarily influenced by taxon and experimental setup (extraction method + scanning mode, *p*-value = 1.97291E-98, ε² = 0.81736339). The highest concentrations were observed in *D. suzukii*, followed by *S. granarius* and *L. japonica*. In the control batch, MU samples yielded higher DNA concentrations than Qi samples across all three taxa. This difference was most pronounced in *D. suzukii* (>4.5-fold), while smaller increases were observed for *S. granarius* (1.23-fold) and *L. japonica* (2.5-fold). MU extracts also exhibited greater variability in DNA concentration compared to Qi extracts (Table S3). When datasets were analyzed separately for each species, the DNA extraction method emerged as the *strongest factor* (Dataset S6). Scanning mode did not significantly affect DNA yield; however, longer irradiation times were associated with reduced DNA concentrations (Fig. S5, Dataset S6).

### DNA fragmentation

The “gentle” extraction methods result in DNA shearing during processing and recover only short fragments (<2,000 bp), precluding assessment of X-ray-induced fragmentation with these approaches. In contrast, proteinase K-based methods (Qi, Xi, MU) enable recovery of long fragments, as demonstrated by the control samples (Dataset S2 and S3). In SE2 series, DNA fragment size was positively associated with DNA concentration and generally decreased with increasing irradiation times (Fig. 8).

**Fig. 8.**
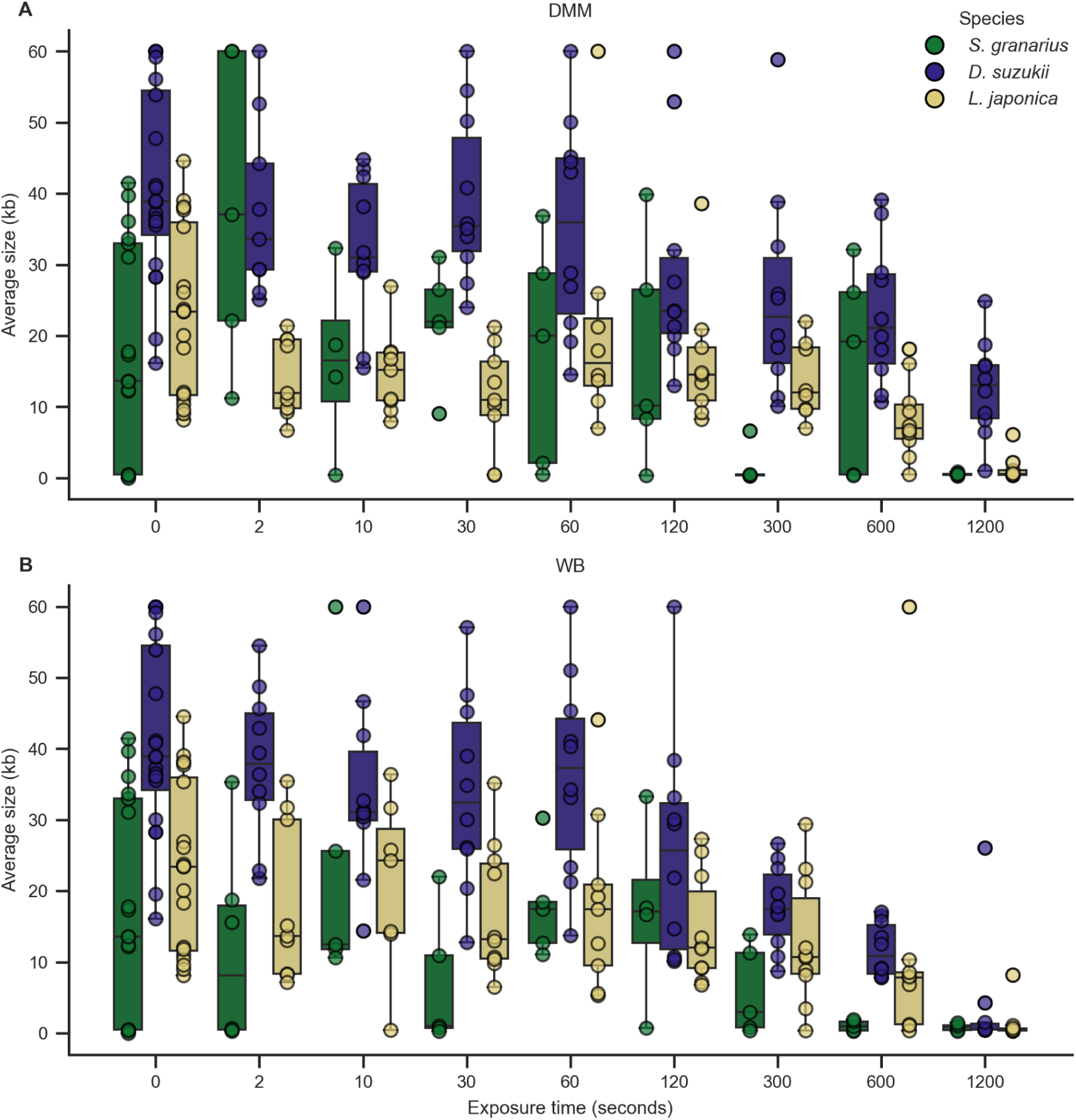
DNA fragmentation. DNA fragmentation across exposure times for two scanning modes: DMM (A), WB (B). Boxplots show the median (central line), interquartile range (boxes), and full range (whiskers) for each treatment at each exposure time. Outliers are indicated as individual points. The two panels compare how the different treatments influence DNA Average size (AS) in samples of *S. granarius, D. suzukii* and *L. japonica* over increasing exposure durations (n = 10).

Across all species, Peak Size (PS) was most strongly affected by exposure time (*p*-value = 1.98731E-27, ε² = 0.281095206,) whereas Average Size (AS) depended on the combination of setup, specifically extraction mode and taxon (*p*-value = 1.38816E-24, ε² = 0.266683321; Dataset S6). When analyzed separately by species, exposure time was the primary factor strongly influencing both PS and AS (Fig. 8, Dataset S6). Generally, for both scanning modes, specimens exhibited fragmentation profiles comparable to controls across all three taxa (Fig. 8, Dataset S2 and S3). Prolonged irradiation resulted in increased fragmentation, with mean AS values of 9,184 bp at 600 seconds and 1,575 bp at 1,200 seconds. For *L. japonica* and *S. granarius,* AS decreased to 6,684 bp (600 seconds) and 523 bp (1,200 seconds). *Drosophila suzukii* samples showed higher DNA concentrations than the other taxa, which was reflected in fragment size distributions (Fig. 8, Dataset S2 and S3). Notably, four *D. suzukii* samples retained limited fragmentation, with AS >15,000 bp at 1,200 seconds.

### DNA barcoding

Exposure time significantly influenced ATQ (*p*-value = 1.51E-05) and CSQ values (*p*-value = 1.01E-07). Additionally, the combination of setup (scanning mode and extraction method) and taxon significantly impacted ATQ and CSQ (p-value = 3.36E-61; 8.51E-33). In *S. granarius* and *L. japonica*, sequencing success rate (Seq), as well as ATQ and CSQ, declined with increasing exposure time (Figs. 9A,B,E,F, S5A,B,E,F and Dataset S6). PCR success rate declined for *S. granarius* under WB and *L. japonica* under DMM and WB. In *D. suzukii* Seq, ATQ, CSQ and PCR success rate seemed virtually unaffected (Figs. 9c,d, and S5C,D).

**Fig. 9.**
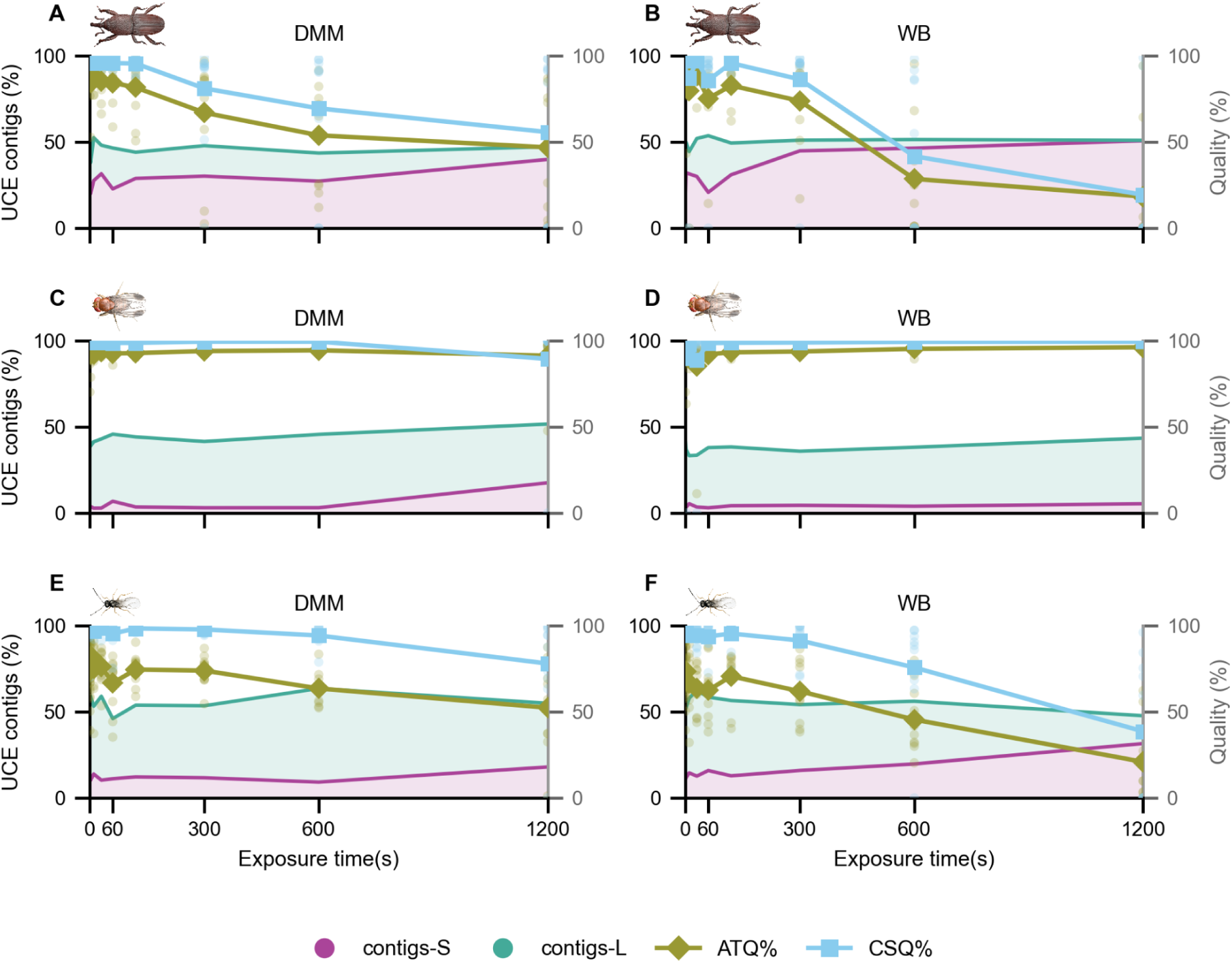
Sequencing quality. Left Y-axis - Scatter plot distribution of overall ATQ% and CSQ% across exposure time range (0 to 1200 seconds; n = 10). Right Y-axis - Quality of UCE sequences - proportion of recovered UCE contigs with length under 1 kb (contigs-S) and over 1 kb (contigs-L) (n = 10). Comparison between scanning mode DMM (left) and WB (right). *S. granarius* (A,B), *D. suzukii* (C,D), *L. japonica* (E,F).

### UCE sequencing

Across all three species, the total bp count was mostly influenced by the exposure time (*p*-value = 0.095250125, ε² = 0.049704537) and combination of setup and taxon (*p*-value = 1.10E-13, ε² = 0.69264616). The total number of contigs was determined by combination of setup and taxon (*p*-value = 2.38E-15, ε² = 0.751064985), while the proportion of loci >1 kb (contigs-L) was impacted by the scanning mode (*p*-value = 0.033501604, ε² = 0.029830174, Dataset S6). Sequence Quality (%) is not affected by exposure time, however, the proportion of loci >1 kb (contigs-L) is decreasing with time for all species and scanning modes, except for *D. suzukii* under WB (Fig. 9).

### Multivariate Analysis (PCA) of species, scanning mode, and exposure time

#### Species separation

Across both scanning modes, species formed distinct clusters along PC1. *D. suzukii* consistently grouped at positive PC1 values, whereas *S. granarius* and *L. japonica* occupied negative PC1 space, indicating that interspecific differences dominate the multivariate structure (Fig. 10).

**Fig. 10.**
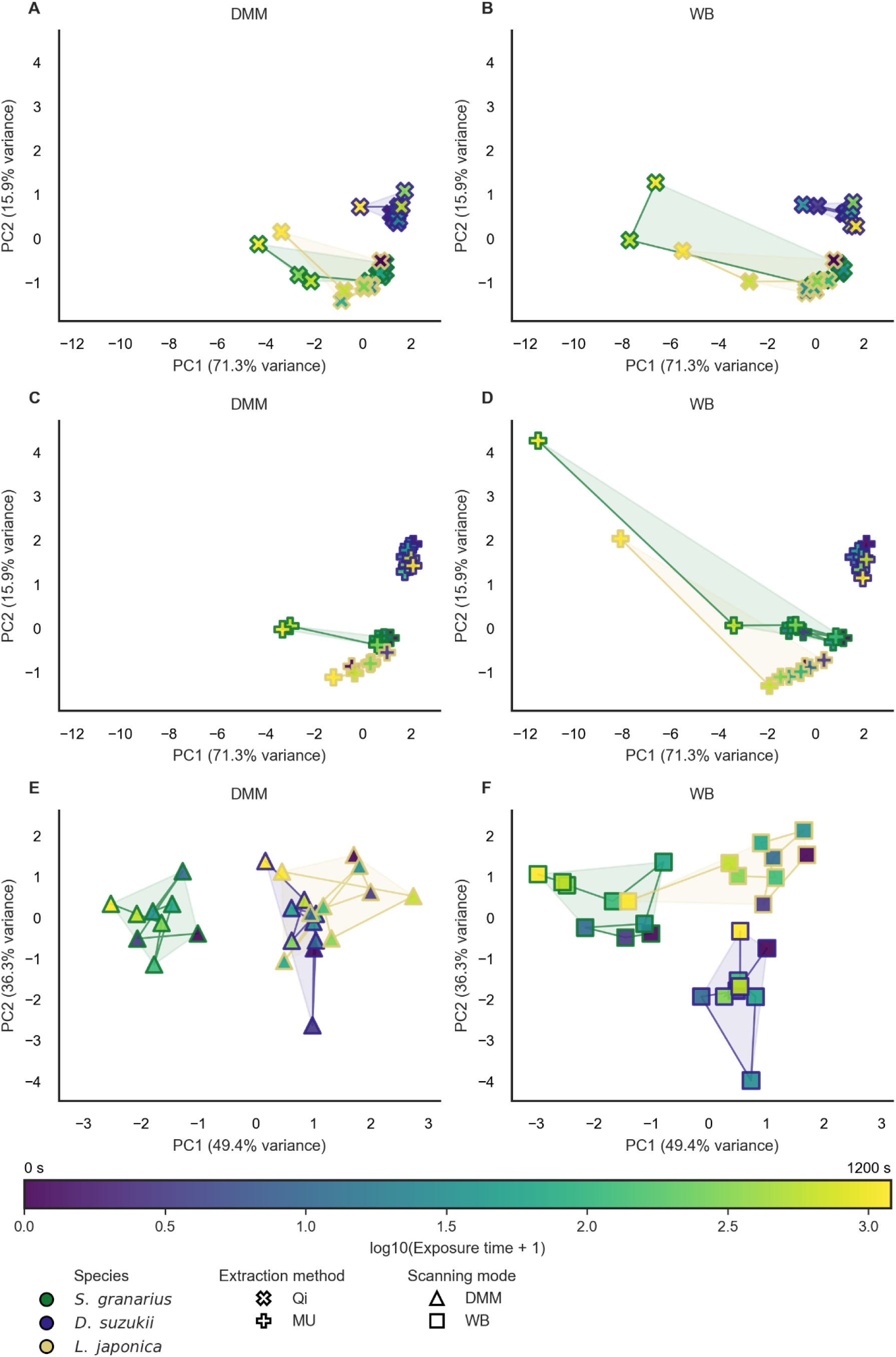
PCA analysis. Combined global principal component analyses (PCAs) of sequencing and assembly metrics across taxa, scanning modes, extraction methods, and exposure times. a-d: Global PCA1 computed using the mean of PCR%, T1Q%, T2Q%, ATQ%, CSQ%, Seq%, and BoxCox% aggregated per Taxon × scanning mode (SM) × extraction method (EM) × exposure time (replicates averaged; n = 5). e,f: Global PCA2 computed UCE data using the mean of contigs_prop, contigs1k_prop, Total-bp, and Count aggregated per Taxon × SM × exposure time (n = 4). Points represent aggregated sample means; outline color indicates taxon/species, marker shape indicates EM in PCA1 and SM in PCA2, and fill color encodes log10(exposure time + 1) (shared color scale; 0-1200 s). Lines connect time-ordered points within each group, and shaded polygons indicate taxon-specific convex hulls within each facet.

#### Scanning mode

Species clustering patterns were largely conserved between DMM and WB, indicating that scanning mode did not fundamentally alter species-level separation. However, WB showed greater dispersion along PC1, suggesting increased exposure-related shifts under WB.

#### Exposure time

In WB longer exposure times shifted samples further along PC1 relative to early exposures, particularly for *S. granarius* and *L. japonica*. This pattern is weaker under DMM.

#### Extraction method

MU vs Qi data points largely overlap within species clusters, indicating minimal separation. This suggests the two methods yield comparable multivariate profiles relative to species and exposure effects.

#### Interaction effects

Species separation is evident on both axes of the PCA (PC1: 49.4%; PC2: 36.3%, Fig. 10E,F), indicating that exposure and species effects are more balanced. Under WB, *D. suzukii* segregated strongly along PC2, highlighting a potential interaction between scanning mode and species at higher exposure levels. Species identity is the dominant determinant of multivariate structure, explaining most variation along PC1. Exposure time introduces secondary, species-specific shifts, particularly under WB scanning. The extraction method has minimal impact relative to biological (species) and technical (scanning mode, exposure time) factors. Overall, WB appears to induce more exposure-related differentiation, whereas DMM yields more compact species clusters.

## Discussion

Our study highlights that high-quality molecular data and high-resolution morphological 3D-data can indeed be obtained from the same insect specimens. However, the order of data acquisition is paramount: SR-µCT scanning must be performed before DNA sequencing, as our results show that all tested DNA extraction protocols are destructive to soft tissues, whereas high quality DNA can still be obtained post-scanning from irradiated specimens. With an optimized, highly automated, high-throughput SR-μCT setup that eliminates the need for in-beam sample alignment, the time required to acquire a high-quality tomogram with 3,000 projections falls well within the “safe zone” for DNA integrity (Figs. 8, 9 and S5). The DNA extraction methods tested are the most established and commonly used in the field of entomology, but the list is not exhaustive. However, since other methods function on the same principles, it would be safe to assume that they would affect the soft tissues in a similar manner.

Both the “gentle” and the proteinase K-based extraction methods resulted in damaged internal structures ranging from increased X-ray absorption to severely or completely digested tissues (Dataset S1). The comparative analysis revealed clear, species-specific responses to the different extraction methods, underscoring that even “gentle” protocols can profoundly alter internal anatomy. QE and HT consistently caused severe shrinkage of soft tissues, most likely due to alkaline-solution induced dehydration, but preserved at least some of the overall organ topology. The proteinase K-based methods produced strong enzymatic degradation in *D. suzukii* and *L. japonica*, with their soft tissues being largely digested, except for the ovaries in *L. japonica* (Figs. 4D, 7F and S3F). This could be attributed to the waxy layer covering the eggshells^48^ of some parasitoid wasps as proteinase K would be ineffective in digesting it. In contrast, *S. granarius* showed very little damage to the soft tissues. This pattern likely reflects differences in cuticle permeability for proteinase K, since weevils have an extremely compact exoskeleton and tight foreclosure of their joints with no exposed arthrodial membranes. The common procedure for DNA extraction of beetles is to open the body cavity, which in this case was not possible since it would defeat the purpose of the study. Additionally, the proteinase K-based methods noticeably changed the internal absorptivity, which is especially striking in Xi specimens (Figs. 4F, 5H and S1H). A potential explanation could be that the Xi (GuSCN-based) lysis buffer infiltrated more readily than proteinase K, leading to retention of sulfur-rich thiocyanate within largely intact tissues, which acted as an internal contrast agent. The apparent lack of damage to the soft tissues is also reflected in the small amounts of DNA recovered for the samples of *S. granarius*. On the other hand, in *D. suzukii* we observed the complete opposite - higher degree of digestion resulted in higher DNA concentration.

Nonetheless, none of the tested methods can be considered truly non-destructive. While the exoskeleton is preserved to some degree, notable morphological damage is present in all specimens post DNA extraction.

In the irradiated samples, DNA yield and quality varied notably between taxa, extraction methods, and irradiation settings. *D. suzukii* consistently produced the highest DNA concentrations across most extractions. The MU method generally outperformed Qi in both concentration and sequencing outcomes but showed greater variability. This likely reflects its purification design, which captures a broader range of fragment sizes - particularly shorter fragments - resulting in higher measured concentrations and improved UCE sequencing success. Although scanning mode did not significantly affect DNA recovery, longer irradiation times consistently reduced DNA concentration. At the same time, DNA integrity was strongly influenced by the extraction method (therefore DNA yield) and exposure time, and related accumulated radiation dose. Proteinase K-based extractions (Qi, MU) recover longer fragments, up to 60,000 bp, making it possible to examine the full fragmentation profile of the samples. In both *S. granarius* and *L. japonica* it is notable that irradiation caused progressive fragmentation, especially at exposures starting at 600 seconds, while *D. suzukii* DNA remained comparatively intact (Fig. 8). This is likely due to much higher DNA yields in *D. suzukii*.

Sequencing success mirrored these trends: in *S. granarius* and *L. japonica*, PCR amplification and sequence quality declined with increased exposure, particularly beyond 300 seconds, while *D. suzukii* remained largely unaffected. MU extracts and the DMM scanning mode produced the best sequencing outcomes. Recovery of UCEs depended primarily on exposure time and extraction method. *Drosophila suzukii* again yielded the most complete and longest loci, while *S. granarius* had the poorest recovery. MU extractions produced slightly longer and more complete loci than Qi. Overall, *D. suzukii* showed the highest DNA stability and sequencing resilience across treatments. Successful sequencing of both DNA barcodes and UCEs could be performed even after more extended irradiation and long fragment size in irradiated samples suggests great potential for long-read sequencing as well.

In our tested setup, DNA fragmentation becomes problematic after approximately 5 minutes of irradiation with both DMM and WB configurations, but the degradation is much more pronounced in the WB samples. A high-quality tomogram with 3,000 projections takes about 60 seconds with the DMM configuration and 25 seconds with the WB configuration. Although WB leads to faster DNA degradation, the much shorter exposure times somewhat compensate for this effect. The PCA results indicate that intrinsic species differences are the dominant determinant of multivariate structure, consistently driving separation along PC1 independent of scanning mode (Fig. 10). Exposure time exerts a secondary, species-dependent effect that is amplified under WB conditions, whereas extraction method contributes comparatively little to overall clustering, suggesting that biological factors and scanning parameters (exposure time) have a stronger influence than scanning mode. Therefore, both modes allow scanning the tested species while preserving sufficient DNA integrity for subsequent DNA barcoding and UCE sequencing. The tomographic image quality obtained with DMM and WB for the same number of projections is highly comparable (Fig. S4). Overall, we conclude that both beam modes perform well and are suitable for combined morphological and molecular analyses. Based on the measured photon flux and the deposited radiation dose, we propose DNA-safe X-ray imaging guidelines transferable to other synchrotron facilities, as similar levels of DNA preservation can be expected at other beamlines if similar photon flux densities, energy distributions, and exposure times are maintained.

In summary, our results demonstrate the practical feasibility of uniting synchrotron X-ray imaging and DNA sequencing within the same insect specimens, thus providing a foundation for integrating genomic and phenomic data at the individual and collection levels. While it is valuable for unique or type specimens, where digitizing both morphology and genetic information is essential, this strategy is particularly promising for high-throughput digitization initiatives. It bridges morphological and molecular data in a reproducible and scalable way, laying the groundwork for a new generation of digital insect collections.

## Material and Methods

### Study specimens

*Drosophila suzukii* and *Leptopilina japonica* were obtained from laboratory strains of Julius Kühn Institute, *Sitophilus granarius* from a laboratory strain of KIT. Before further processing, fresh individuals were fixed in 99% ethanol and transferred into 0.2 ml microtubes without additional dissection.

### Experimental concept

To systematically evaluate the compatibility of synchrotron X-ray imaging and DNA sequencing within the same insect specimens, we designed two complementary experiments (Fig. 2).

#### Experiment 1: DNA extraction-induced damage on morphology

Five commonly used DNA extraction protocols were evaluated. To directly assess extraction-induced damage, the same individual specimens were scanned using high-throughput synchrotron microtomography (Synchrotron Experiment 1) both before and after DNA extraction. Tomographic imaging was performed using a monochromatic beam (DMM), which allows precise control of photon energy and deposited dose. Each specimen was scanned with either 1,500 projections, 3,000 projections (about the recommended number for given setup^49^), or 6,000 projections. For comparison with another beam mode, additional tomograms were acquired using a polychromatic white beam (WB) with the same projection numbers. Following the synchrotron experiment, DNA was extracted from the scanned specimens. Two non-proteinase K-based (“gentle”) DNA extraction methods were applied: HotSHOT (n = 45) and QuickExtract (n = 45), as well as three proteinase K-based methods: an in-house protocol employing a Xiril Automated Station (n = 45) and two Qiagen kits - DNeasy Blood & Tissue (n = 45) and QIAamp Micro (n = 15).

#### Experiment 2: X-ray-induced DNA degradation

Specimens were irradiated for durations ranging from 2 to 1,200 s under DMM and WB (Synchrotron Experiment 2) in order to quantify DNA fragmentation as a function of irradiation time and beam parameters. Photon flux was measured experimentally, and the absorbed radiation dose was estimated by numerical dose simulations. Subsequent DNA extraction was performed with two proteinase K-based methods with Qiagen kits - DNeasy Blood & Tissue (n = 240) and QIAamp Micro (n = 240).

### Synchrotron experiment 1 (SE1): High-throughput X-ray microtomography

Samples (Dataset S1) were scanned at the IMAGE beamline^50^ of the KIT Light Source using the UFO-I high-throughput X-ray microtomography setup equipped with a robotic sample exchanger. We used a fast indirect detector system consisting of a 25 µm LSO scintillator^51^, a double objective white beam microscope^52^ (Optique Peter) and a 12 bit pco.dimax high-speed camera (Excelitas PCO GmbH) with 2,016 x 2,016 pixels. A 5x magnification resulted in an effective pixel size of 2.44 µm.

195 individuals (65 of each species) were scanned using a monochromatic X-ray beam diffracted by a double multilayer monochromator. A superconducting wiggler with a magnetic field of 2.7 T provided a maximum flux density at 15.2 keV with an energy bandwidth of ∼3% (Fig. S6). To reduce the heat load on the monochromator, the beam was pre-filtered with 3 mm pyrolytic graphite. From each species, 20 specimens were scanned using 1,500 projections, 25 using 3,000 projections, and 20 using 6,000 projections, resulting in exposure times of 39.5 s, 78 s, and 156 s, respectively. In each case, 100 dark-field images and 200 flat-field images were acquired for background correction. After DNA extraction, all individuals scanned in DMM mode were re-scanned using the same setup.

195 samples (65 of each species) were scanned with a filtered polychromatic beam (“white beam”, WB), by filtering the spectrum of the superconducting wiggler (corresponding to a magnetic field of 1 T) with 10 mm pyrolytic graphite sheets. The resulting spectrum, peaked at approximately 16.5 keV and having a full width at half maximum of approximately 11 keV (Fig. S6), is free from the low energy components. Exposure times during tomography were 12.5 s (1,500 projections), 25 s (3,000), and 50 s (6,000). The total exposure times matched those used in DMM mode.

Like in DMM mode, 20 specimens from each species were scanned using 1,500 projections, 25 using 3,000 projections, and 20 using 6,000 projections. For each scan, 100 dark-field images and 200 flat-field images were acquired. In contrast to the DMM series, the WB samples were not re-scanned.

Online and final data processing included dark and flat field correction and phase retrieval of the projections based on the transport of intensity equation^53^. Online tomographic reconstruction was done with the UFO framework^54^. The final 3D tomographic reconstruction was performed by tofu^55^ and included ring removal, 8-bit conversion and blending of the phase and absorption 3D reconstructions.

#### Post-processing of tomographic data

For each species and extraction method, we selected a representative pair of tomographic volumes based on scans of the same individual taken before and after DNA extraction. The blended 8-bit versions of the data were aligned, cropped and isolated from the background with Amira 3D 2022.2. The processed volumes were imported into Drishti 2.5.1^56^, digitally sectioned along the center of the thorax and rendered. In Adobe Photoshop, contrast and brightness of the renderings were adjusted.

For each species, a representative surface model was created based on a blended 8-bit tomogram of a pre-extraction scan. Amira 3D 2022.2 was used to pre-segment the exoskeleton and selected soft tissues. The pre-segmented labels served as input for semi-automatic segmentation with Biomedisa^57^. The Biomedisa results were re-imported into Amira and minor errors were corrected. The final label fields were converted into polygon meshes, which were exported as OBJ files and reassembled and smoothed in Cinema 4D 2026, which was also used for artificial coloring, virtual sectioning and final surface rendering.

### Synchrotron experiment 2 (SE2): Irradiation series

480 samples were irradiated for different durations (Dataset S2) at the IMAGE beamline without tomographic imaging. Using a monochromatic X-ray beam with the parameters described above for DMM mode, 10 samples of each species were irradiated for 2, 10, 30, 60, 120, 300, 600 and 1,200 seconds. An identical series was irradiated with a polychromatic X-ray beam with the parameters described above for WB mode. For both series, the samples were placed on the sample stage by the beamline’s robotic sample exchanger and irradiated for the desired duration. The opening and closing times of the beam shutter were recorded. Full-body exposure was ensured by monitoring the samples with the X-ray detector during irradiation.

#### Measurements of X-ray photon flux density at sample position

The experimental setup used for measuring the incident X-ray photon flux at the sample position included: a) a silicon photodiode (Hamamatsu S3590-09) with a thickness of 300 µm and coated with a 200 nm SiO₂ protective layer, b) a low-noise current preamplifier (DLPCA-200, FEMTO Messtechnik GmbH), which amplifies the generated photocurrent, and c) a four channel ultra-high linearity Voltage-to-Frequency converter (Model N101VTF NOVA R & D, Inc), to convert the photocurrent into a frequency signal.

Before proceeding with the quantitative estimation of the flux density, we assessed the detector response under varying beam intensities, by recording the total photocurrent as a function of filter transmission (3 mm to 10 mm pyrolytic graphite and 3 mm pyrolytic graphite + 1 mm silicon carbide). Results proved a linear correlation between the measured photocurrent and the nominal transmission values of the filters (Fig. S7). This outcome enabled the use of the diode for quantitative measurements of X-ray flux regime.

The quantitative estimation of the impinging X-ray photon flux has been done taking into account the diode geometry, the energy-dependent absorption within the active layer, and transmission losses due to any entrance window or passive layers. In detail, the X-ray photon flux density is given from the following expression^58^:

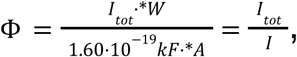

where

– I_tot_ is the total photocurrent generated in the diode;
– W is the mean energy required to produce one electron-hole pair (3.68+/- 0.02) eV;
– k is the photon energy, equal to 15.2 keV in this specific case;
– F is the energy fraction deposited in the Si diode (thickness = 300 µm), equal to 0.48 at the 15.2 keV energy used for the experiment;
– A is the irradiated area of the diode;
– I is the photocurrent produced by one incident X-ray photon.

In this estimation, the absorption within the front oxide dead layer (∼200 nm thick) and the effect of radiation backscattered from the backing material were neglected. To define the geometry of the incident beam on the photodiode and to align it accurately, a pair of motorized slits was used. The beam aperture of 0.85 mm x 0.930 mm (H x V) was visualized with the indirect detector used to acquire the tomographic scans and described for SE1.

As for the total uncertainty in the flux calculation (∼ 5 %), this was estimated by calculating the maximum semi dispersion between the flux density calculations. The calculated value of the photon flux density at the DMM energy of 15.2 keV was equal to 1.17*10^12^ ph/mm^2^/s at a ring current of 125 mA and under a prefiltering of 3 mm pyrolytic graphite.

#### Simulation of the deposited dose

To estimate the deposited dose in the specimen, a Geant4^59,60,61^ (version 11.3.2) based Monte-Carlo dose simulation was developed. In this simulation, samples composed of a cylindrical plastic tube filled with 100% ethanol and three different insect shapes were modeled. The insect shapes resemble coarse estimations of *S. granarius*, *D. suzukii* and *L. japonica* (Fig. S8). The “insects” were represented by a composite material consisting of approximately 70 % dry biomass and 30 % of absolute ethanol by mass. The dry biomass was modeled with a simplified composition of 50% carbon, 7% hydrogen, 9% nitrogen, 33% oxygen, and 0.5 % each of phosphorus and sulfur, based on whole-body elemental data across multiple insect taxa^61,63^; and the dominant biochemical components chitin, proteins, and lipids^64^. Ethanol contributes additional carbon and hydrogen while reducing the overall oxygen fraction compared to hydrated tissue. The bulk density of the ethanol-preserved insect material was estimated at approximately 0.95 g/cm³, based on a weighted combination of dry tissue (∼1.3–1.4 g/cm³) and 100% ethanol (0.789 g/cm³). The specimen, sample tube and ethanol were then exposed to 10,000,000 photons with an energy of 15.2 keV and the absorbed dose was sampled in each of the different volumes. By assuming the photon flux density of 9.37 * 10^11 ph/mm²/s the dose rate was calculated. For the filtered white beam exposure the simulated spectrum was applied in the dose simulation resulting in these dose rates (Table S1). We assume the error of the dose simulation to be 10% and with the error of 5% of the flux measurement/simulation we can assume an error of 11.2% for the dose rate. For the total dose an error of 0.1s in the exposure time is taken into account. All values were normalized to a ring current of 125 mA that is close to the average current during our experiments.

### Extraction methods

#### “Gentle” extraction methods

DNA leaching with HotSHOT (HT) does not involve muscle digestion with proteinase enzymes and is therefore the least destructive approach. The specimens were immersed in 15 µl HotSHOT Alkaline buffer (25 mM NaOH, 0.2 mM Na2EDTA, pH12) and incubated for 20 minutes at 65 °C (soft-bodied insects: *D. suzukii*)^65^ or for 30 minutes at 95 °C (heavily sclerotised insects: *L. japonica* and *S. granarius*)^66^. After incubation, 15 µl of Neutralising buffer (40 mM Tris-HCl, pH5) was added. The extracts were transferred to 1.5 µl vials and purified using the purification membrane columns from the DNeasy Blood & Tissue Kit from Qiagen. The elution was performed in two steps: 20 µl + 15 µl elution buffer, each with 15 minute incubation at room temperature. All HotSHOT extracts were then concentrated to 10 µl.

A commercial version of the “gentle” extraction buffer was used for comparison. The QuickExtract (QE) buffer from Lucigen was used in a similar manner: 25 µl with a 5 minute incubation at 95 °C followed by 15 minutes incubation at 65 °C. The extracts were transferred to 1.5 µl vials and purified using the purification membrane columns from the DNeasy Blood & Tissue Kit from Qiagen, as described above. All QE extracts were then concentrated to 10 µl.

#### Protein K-based extraction methods

The most destructive DNA extraction protocol tested here uses a 30 h incubation with proteinase K at 56 °C. We employed a Xiril Automated Station (Xi) for DNA purification^67,68^, elution was performed in 60 µl 1x TE buffer. For comparison, commercial alternatives were also tested: the DNeasy Blood & Tissue Kit from Qiagen (Qi), following the manufacturer’s protocol with additional modifications^69^. Elution was performed in two steps, with 50 µl elution buffer each, resulting in a total volume of 100 µl. A third proteinase K-based method was employed, specifically designed for the recovery of fragmented DNA, such as from old museum specimens (MU). The Qiagen QIAamp Micro DNA extraction kit was used following the manufacturer’s protocol with adjustments^70^, except for the grinding of specimens. Elution was performed in two steps, with 20 µl elution buffer each, to maximize yield, resulting in a total volume of 40 µl.

#### Negative controls

All extraction methods included a blank sample to monitor reagent contamination. In addition, non-irradiated specimens of each species were included as negative controls (SE1: two specimens each; SE2: 10 specimens each).

### Quantification of DNA extracts

The quality check of the extracted DNA was three-fold: (1) we quantified the DNA concentration in the extracts using the High-Sensitivity Assay for double stranded DNA using a Qubit fluorometer (1 µl of DNA extract for HT and QE; 2 µl of DNA extract for Qi, Xi and MU). (2) Fragmentation of the DNA was documented through capillary electrophoresis on Fragment Analyser (FA) Version 1.2.0.11 (Agilent Technologies, USA) with the HS Genomic DNA 50kb Kit (DNF-468-0500). Sample preparation followed the manufacturer’s protocol provided with the kit. In addition, samples were treated with 1 µl of Ribonuclease A (90 U/mg, watery solution of 10 mg/ml), followed by a 10 minute incubation at room temperature prior to capillary injection. (3) Samples were tested for sequencing in a standard DNA barcoding approach and the recovery of Ultra Conserved Elements (UCE).

#### Barcoding and sequencing

PCR amplification was performed in a 25 µl reaction with 10 µl DNA template (HT, QE) or 4 µl DNA template (Qi, Xi, MU). The COI fragment was targeted with the LCO1495 - HCO2198 primers^71^. PCR success was assessed through agarose gel electrophoresis with the Bio-Rad Gel Doc XR+ System. Amplicons were sequenced bidirectionally through Sanger sequencing, raw reads were assembled and proof read in Geneious Prime. Final consensus sequences were checked for contamination through BLAST against the BOLD database^72^.

#### UCE library preparation and sequencing

Ultraconserved Elements (UCEs) were extracted and used as molecular markers due to their broad applicability in resolving phylogenetic relationships and population genomics analyses^19^. Our analyses include a total of 117 samples, with 39 samples of each species. After extraction, 50–100 ng of DNA were sheared by sonication (Qsonica) to an average fragment size of ∼ 450–600 bp and used for the library preparation (Kapa Hyper Prep Library Kit, Kapa Biosystems) incorporating “with-bead” (SeaPure) cleaning steps^73^. Libraries were tagged with unique combinations of iTru adapters^74,75^, and posteriorly pooled for target enrichment using the Coleoptera 1.1Kv1, Diptera 2.7Kv1, and Hymenoptera 2.5Kv2P Microarray probe sets, which target 1,172, 2,711, and 2,590 UCE loci from Coleoptera, Diptera, and Hymenoptera, respectively. We combined samples into an equimolar pool and sequenced them at the Biomarker Technologies (BMK) GmbH using an Illumina Nova PE150 platform. After sequencing, the PHYLUCE v1.7.1 protocol was used for processing UCE in preparation for phylogenomic analyses^75^. After quality control and cleaning, samples were assembled using SPAdes^76^. Assembled contigs were matched to the UCE probes, and calculated statistics on UCE assemblies for individual taxa.

### Data Analysis

#### DNA concentration

To identify potential data outliers, a Z-score analysis was performed on the DNA concentration data (Dataset S1 and S2). Prior to further analysis, the Box-Cox transformation^77^ was implemented on the data. The changes in DNA concentration in relation to exposure time and the comparison between the two scanning modes were assessed on scatter plots for each species and extraction method separately.

#### DNA fragmentation

DNA fragmentation levels were assessed based on Peak Size (PS) and Average Fragment Size (AS). PS and AS values were exported from the electropherograms (Dataset S3) and used for follow up analyses (Dataset S2). To identify potential data outliers, a Z-score analysis was performed on the DNA fragmentation data. Distribution and variance of AS values were assessed via boxplots.

#### DNA Barcoding

PCR amplification success rate was encoded as follows: 0 = fail and 1 = passed, based on gel inspection. Consensus sequences were assembled *de novo*, trimmed and checked via visual inspection of the contigs *vs* electropherograms in Geneious Prime. Raw trace quality data and Consensus sequence quality (CSQ) were exported in batch from Geneious Prime. Average trace quality (ATQ) was calculated based on raw reads for each sample individually. For samples that failed amplification and/or sequencing (contig could not be produced), CSQ was encoded as 0. Sequencing success rate was quantified as the sequencing of the target organism and encoded as follows: 0 = fail, i.e. contamination OR sequence could not be assembled due to failed amplification and/or sequencing, and 1 = passed (Dataset S2). ATQ, CSQ, PCR and Sequencing success rates were plotted on area and line charts individually per species, data was aggregated to mean or median values according to parametric/non-parametric distribution.

#### UCE sequencing

To assess the success of UCE sequencing we evaluated sequencing Quality (%) - the proportion of recovered UCE loci, below (contigs-S) and over (contigs-L) 1k bp, the Total bp count and the length of the contigs (Dataset S4). UCE sequencing Quality (%) was plotted as area charts individually per species.

#### Statistical analysis

Statistical analysis was performed on the following data: DNA concentration (Box-Cox transformed data), PS and AS, PCR and Sequencing success rate, ATQ, CSQ, Total bp count, contigs-S, contigs-L, and the length values of the recovered loci (min, median, mean, max and 95 Cl length; Dataset S5). To assess data distribution, the Shapiro-Wilk test was employed to determine whether datasets met parametric or non-parametric assumptions^78^. Levene’s Test was run to check for Homogeneity of Variance^79^. One-Way ANOVA was run for parametric data. Kruskal-Wallis was used for non-parametric data^80^. Fisher exact was employed for binary data. Post-hoc statistical analyses were performed if ANOVA and Kruskal-Wallis were significant: Tukey’s HSD^81^ and Dunn’s test^82^, respectively. Eta-squared (η²) and Epsilon-squared (ε²) were used to determine the effect size for the parametric and non-parametric cases, respectively. Principal Component Analysis (PCA) was performed on standardized metrics to reduce dimensionality and visualize patterns among samples. The first two principal components, capturing the largest variance, were used to inspect clustering according to species, scanning mode, extraction method, and exposure time. Multivariate inference (PERMANOVA/ Freedman-Lane permutation tests and PERMDISP-like dispersion tests) was performed on Euclidean distances of the standardized feature matrices used for each PCA.

Analyses were performed in python using Scikit-learn^83^, SciPy^84^ and Scikit-bio^85^. Graphs and figures were created in python using Matplotlib^86^ and Seaborn^87^.

## Data availability

All processed tomograms analyzed in this study are accessible through the RADAR4KIT repository (https://radar.kit.edu/radar/en/), (access link will be activated upon publication): https://dx.doi.org/10.35097/qj9139epc345z671 (*S. granarius* (QE)), https://dx.doi.org/10.35097/bx1n6qb1uncr4tbz (*S. granarius* (HT)), https://dx.doi.org/10.35097/zay570gp7tz44jaz (*S. granarius* (Qi)), https://dx.doi.org/10.35097/nudnh7vhkyy9nwsz (*S. granarius* (Xi)), https://dx.doi.org/10.35097/r6uc0zba0tacjra5 (*S. granarius* (MU)), https://dx.doi.org/10.35097/unzhj56w00kpe6sn (*D. suzukii* (QE)), https://dx.doi.org/10.35097/fjk3tkdpzxu38gka (*D. suzukii* (HT)), https://dx.doi.org/10.35097/f78vws3jn7a6xzq0 (*D. suzukii* (Qi)), https://dx.doi.org/10.35097/su4qac3u4g47mad1 (*D. suzukii* (Xi)), https://dx.doi.org/10.35097/0w60bkscu5hdy3gt (*D. suzukii* (MU)), https://dx.doi.org/10.35097/3m4h8m2q5c404ttp (*L. japonica* (QE)), https://dx.doi.org/10.35097/5cy5e8dtxbyb949g (*L. japonica* (HT)), https://dx.doi.org/10.35097/mq1n0wbqkmmqvw4m (*L. japonica* (Qi)), https://dx.doi.org/10.35097/c0qaa08vbgvmbnp8 (*L. japonica* (Xi)), https://dx.doi.org/10.35097/pmb9491agv5aene6 (*L. japonica* (MU))

Temporary access links during review: https://radar.kit.edu/radar/en/dataset/qj9139epc345z671?token=MPfexzAdwmdKZkNFanUV https://radar.kit.edu/radar/en/dataset/bx1n6qb1uncr4tbz?token=WbDGJkqTczMOlClKdRug https://radar.kit.edu/radar/en/dataset/zay570gp7tz44jaz?token=LvAOEnXlpfANXVTsZoaz https://radar.kit.edu/radar/en/dataset/nudnh7vhkyy9nwsz?token=RVJEyDrqEXzQLRZeFjTS https://radar.kit.edu/radar/en/dataset/r6uc0zba0tacjra5?token=vEbysXHtozNscOXeelvi https://radar.kit.edu/radar/en/dataset/unzhj56w00kpe6sn?token=XFSSmawpRKKJHxiDFPHv https://radar.kit.edu/radar/en/dataset/fjk3tkdpzxu38gka?token=foeWJFZtQRCAgSiQhkCd https://radar.kit.edu/radar/en/dataset/f78vws3jn7a6xzq0?token=TrYZjxtoIugoqAAUmqKr https://radar.kit.edu/radar/en/dataset/su4qac3u4g47mad1?token=UzQqrGiwYFhLlfQtJyDF https://radar.kit.edu/radar/en/dataset/0w60bkscu5hdy3gt?token=VaCzlOVfVAddxUoAZhTJ https://radar.kit.edu/radar/en/dataset/3m4h8m2q5c404ttp?token=OoQYkCUOiXUioTndjLBB https://radar.kit.edu/radar/en/dataset/5cy5e8dtxbyb949g?token=xvaPApZRTkETeYscTZdQ https://radar.kit.edu/radar/en/dataset/mq1n0wbqkmmqvw4m?token=wkHGjgpvOaWLExtYBNjk https://radar.kit.edu/radar/en/dataset/c0qaa08vbgvmbnp8?token=jWuPCJxwmcKRfocfxlzE https://radar.kit.edu/radar/en/dataset/pmb9491agv5aene6?token=dMVTmNPkUjIUuifFCkeD

DNA barcode and BLAST data, as well as UCE sequences are accessible through the RADAR4KIT repository (https://radar.kit.edu/radar/en/) (access link will be activated upon publication): https://dx.doi.org/10.35097/wdv44as289jjcd2v. Temporary access link during review: https://radar.kit.edu/radar/en/dataset/wdv44as289jjcd2v?token=rNUlcEAAHxiGkdgHTCMp

The source code of the dose simulation is accessible under https://doi.org/10.5281/zenodo.17864742.

## Acknowledgements

Research at KIT was supported by the project SMART-Morph (05K2022) via the German Federal Ministry of Research, Technology and Space (BMFTR). We gratefully acknowledge the data storage service SDS@hd supported by the Ministry of Science, Research and the Arts Baden-Württemberg (MWK) and the German Research Foundation (DFG) through grant INST 35/1503-1 FUGG. We acknowledge the KIT Light Source for provision of instruments at their beamlines and we would like to thank the Institute for Beam Physics and Technology (IBPT) for the operation of the storage ring, the Karlsruhe Research Accelerator (KARA).

## Author contributions

C.L.-V., A.R. & T.v.d.K. designed the study. A.H., J.M. & T.v.d.K. prepared samples. A.C., E.H., J.H., J.O., P.P., R.S., M.Z. & T.v.d.K. performed X-ray experiments. C.L.-V. & D.M.-R. performed DNA extraction and sequencing. C.T. & T.F. reconstructed tomographic data. C.L.-V., C.T., T.F. & T.v.d.K. analyzed tomographic data. A.C. & M.Z. calculated the deposited radiation dose. C.L.-V., A.R., D.M.-R. & C.R. analyzed DNA fragmentation and sequencing results. C.R., T.B., L.K. & T.v.d.K. supervised the work. C.L.-V. & T.v.d.K. drafted the manuscript. All authors contributed to writing the manuscript.

## Supplementary information

**Fig. S1.**
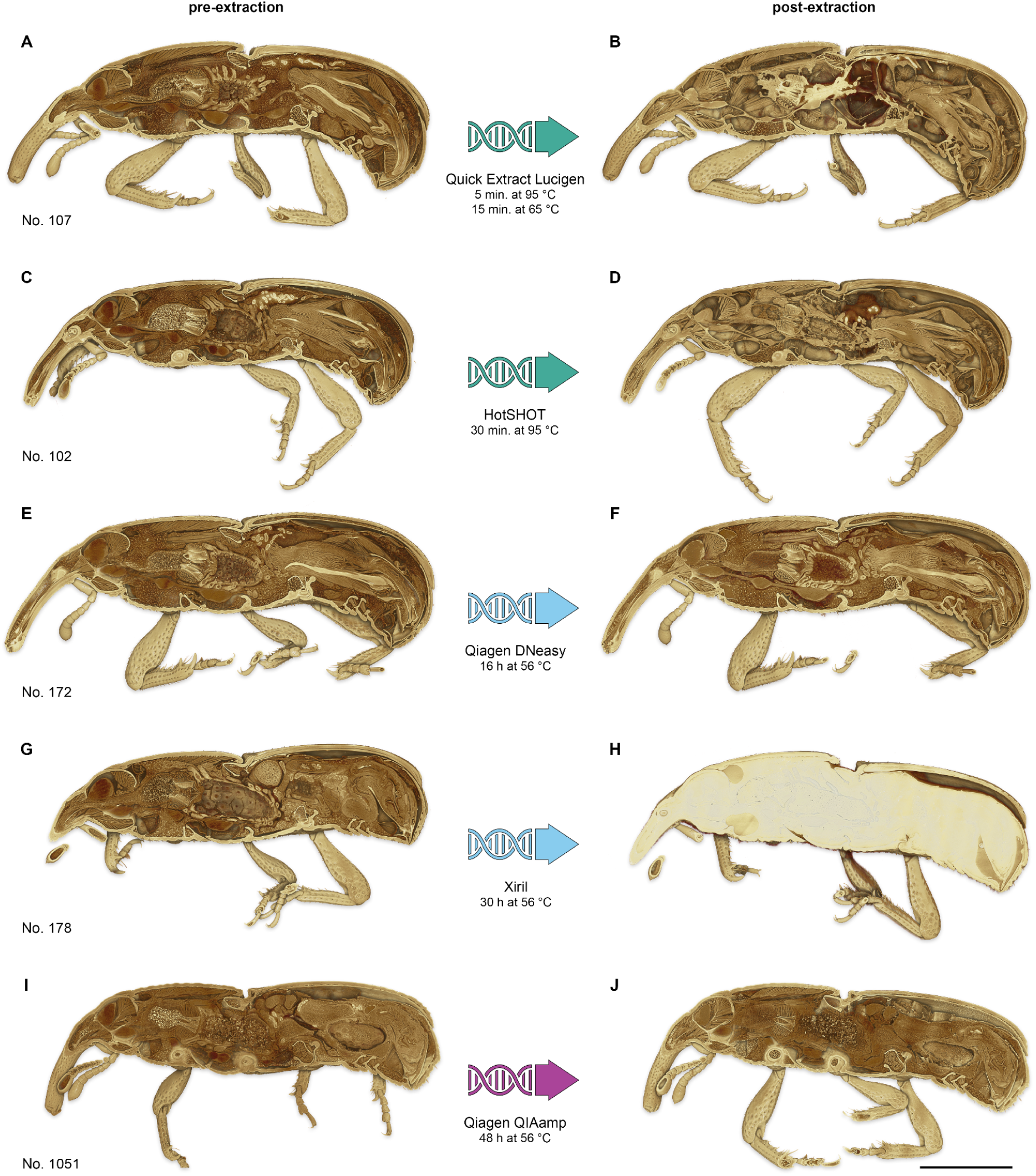
Sectioned volume renderings of *Sitophilus granarius* before and after DNA extraction. Examples show the same individuals scanned pre- and post-extraction highlighting the effects of different DNA extraction methods on internal anatomy. Scale bar = 1 mm.

**Fig. S2.**
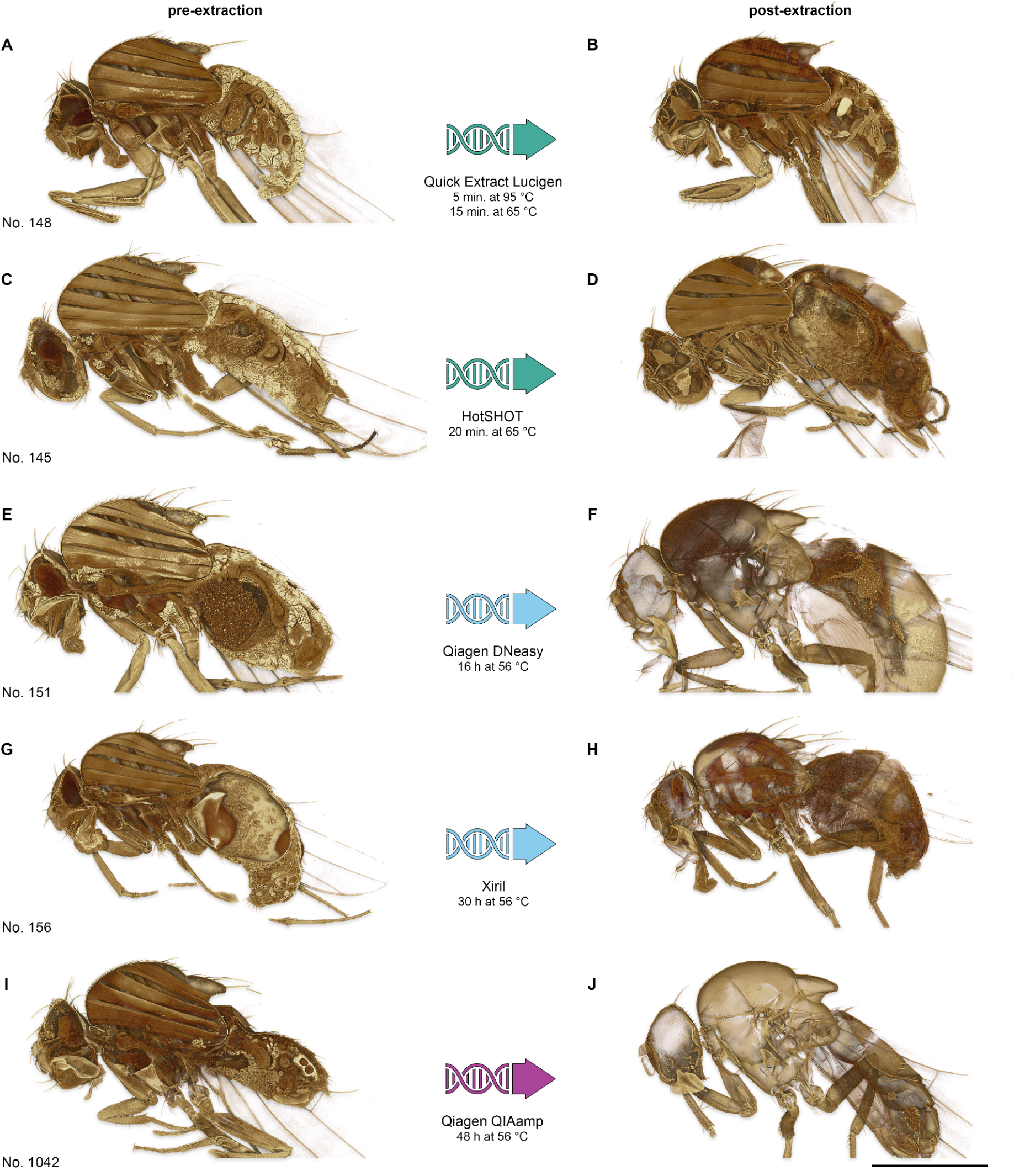
Sectioned volume renderings of *Drosophila suzukii* before and after DNA extraction. Examples show the same individuals scanned pre- and post-extraction highlighting the effects of different DNA extraction methods on internal anatomy. Scale bar = 1 mm.

**Fig. S3.**
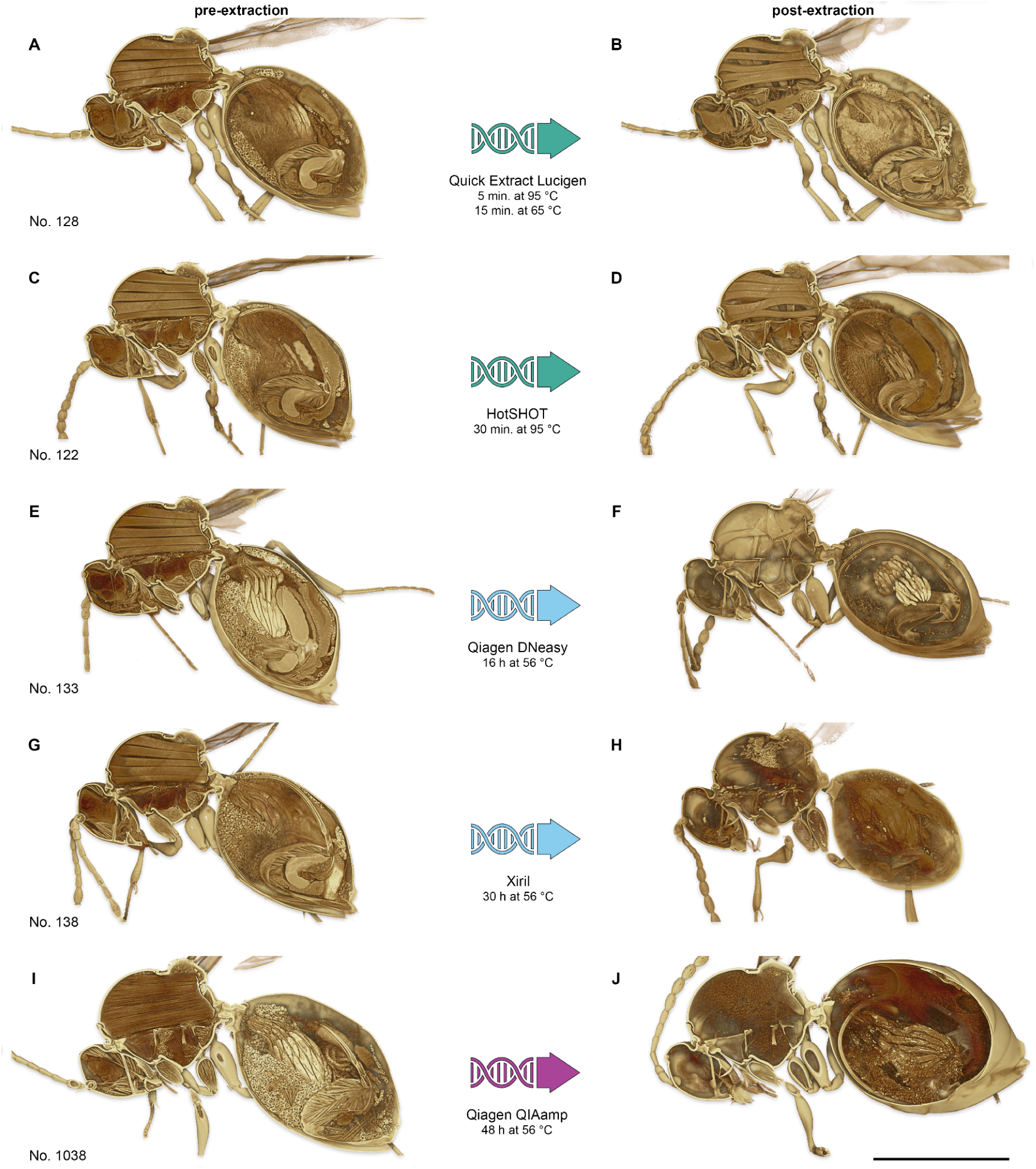
Sectioned volume renderings of *Leptopilina japonica* before and after DNA extraction. Examples show the same individuals scanned pre- and post-extraction highlighting the effects of different DNA extraction methods on internal anatomy. Scale bar = 1 mm.

**Fig. S4.**
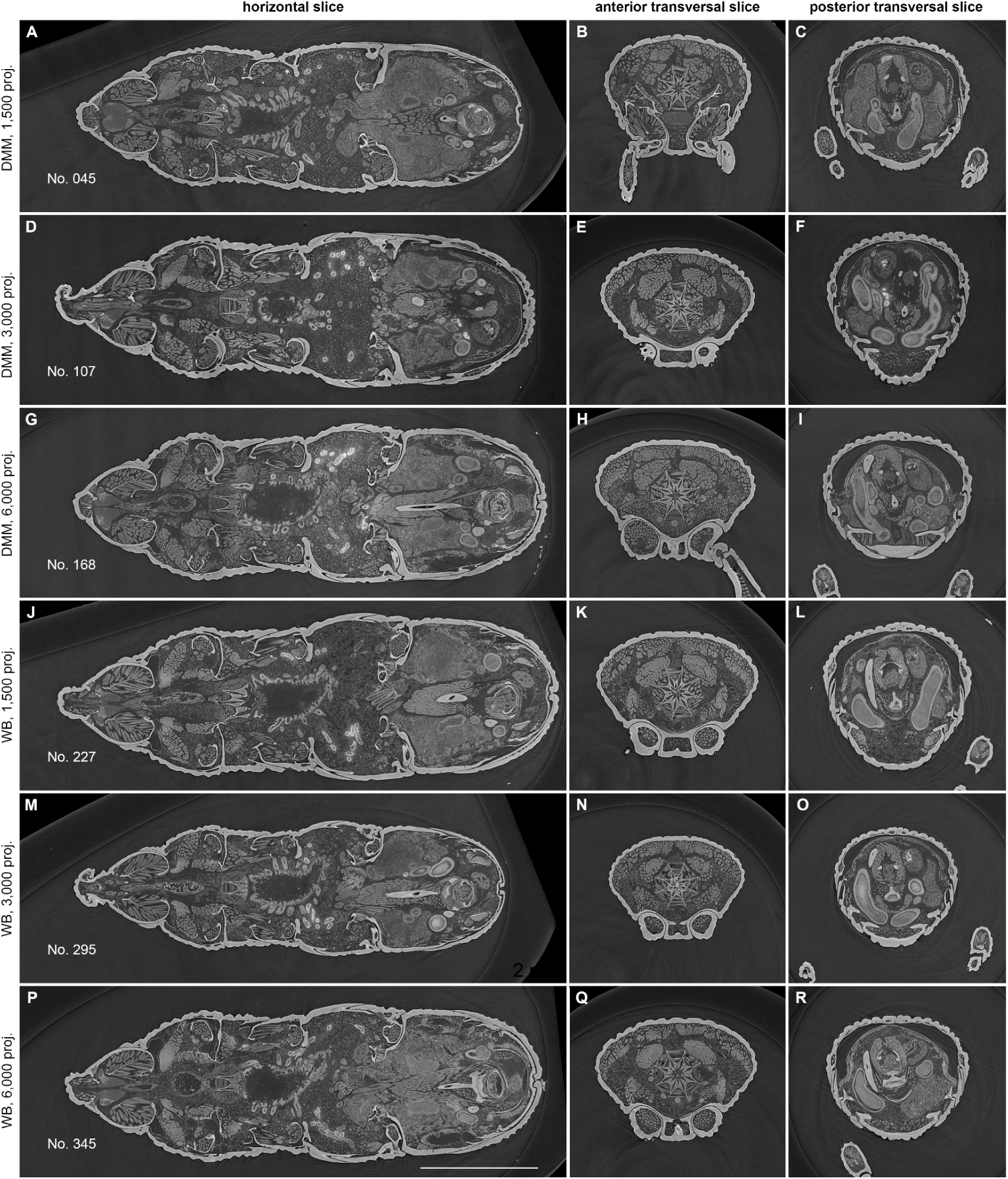
Comparison of tomographic data acquired with different scanning modes. Representative slices of *Sitophilus granarius* from tomograms based on 1,500, 3,000 and 6,000 projections (proj.), acquired with polychromatic beam (DMM) and white beam (WB). Scale bar = 1 mm.

**Fig. S5.**
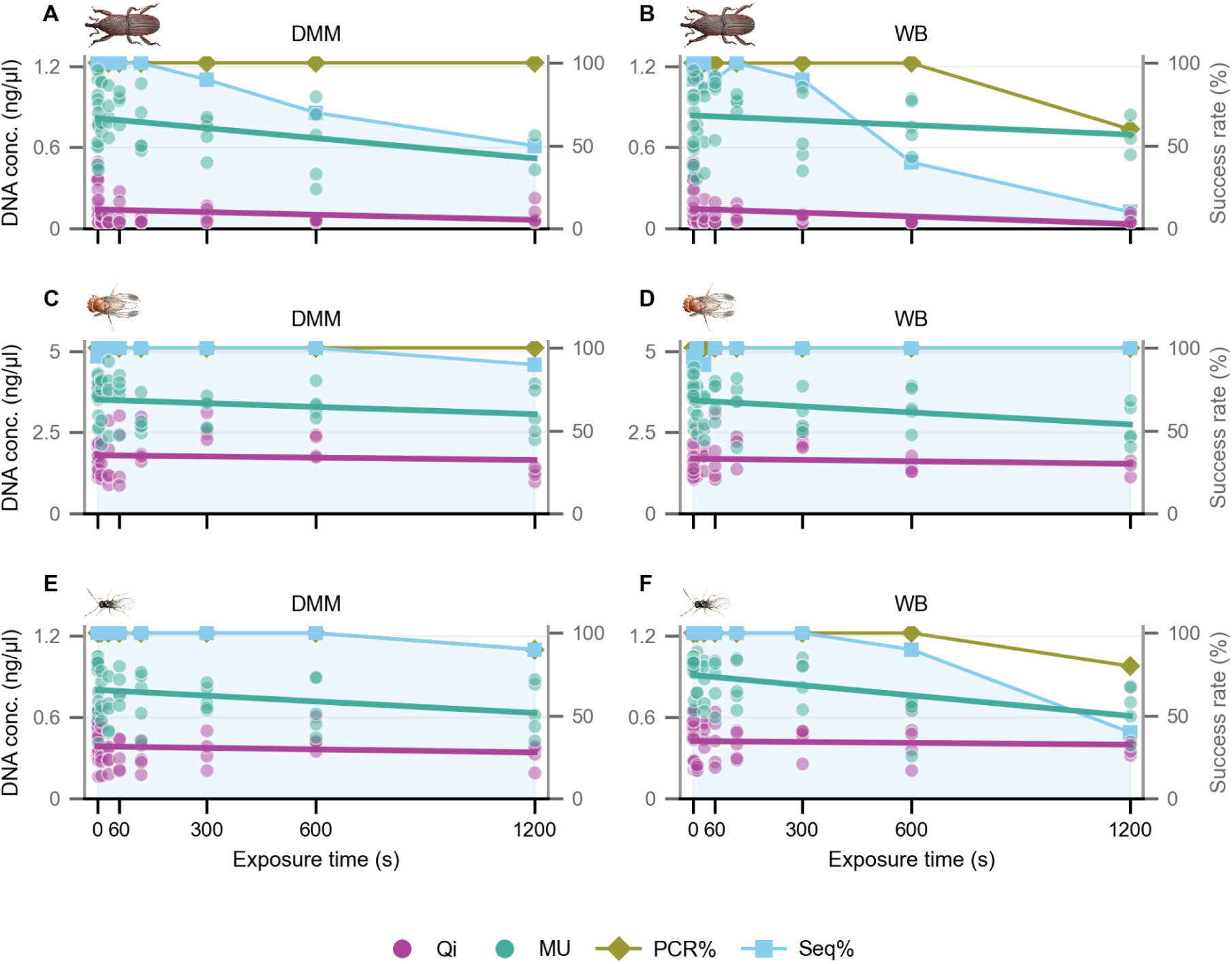
DNA quality. Left Y-axis - Scatter plot distribution of DNA concentration values (BoxCox transformed data) across exposure time range (0 to 1200 seconds) with linear trend lines (n = 5). Right Y-axis - Overall PCR and Sequencing success rates (n = 10). Comparison between scanning mode DMM (left) and WB (right). *S. granarius* (A,B), *D. suzukii* (C,D), *L. japonica* (E,F).

**Fig. S6.**
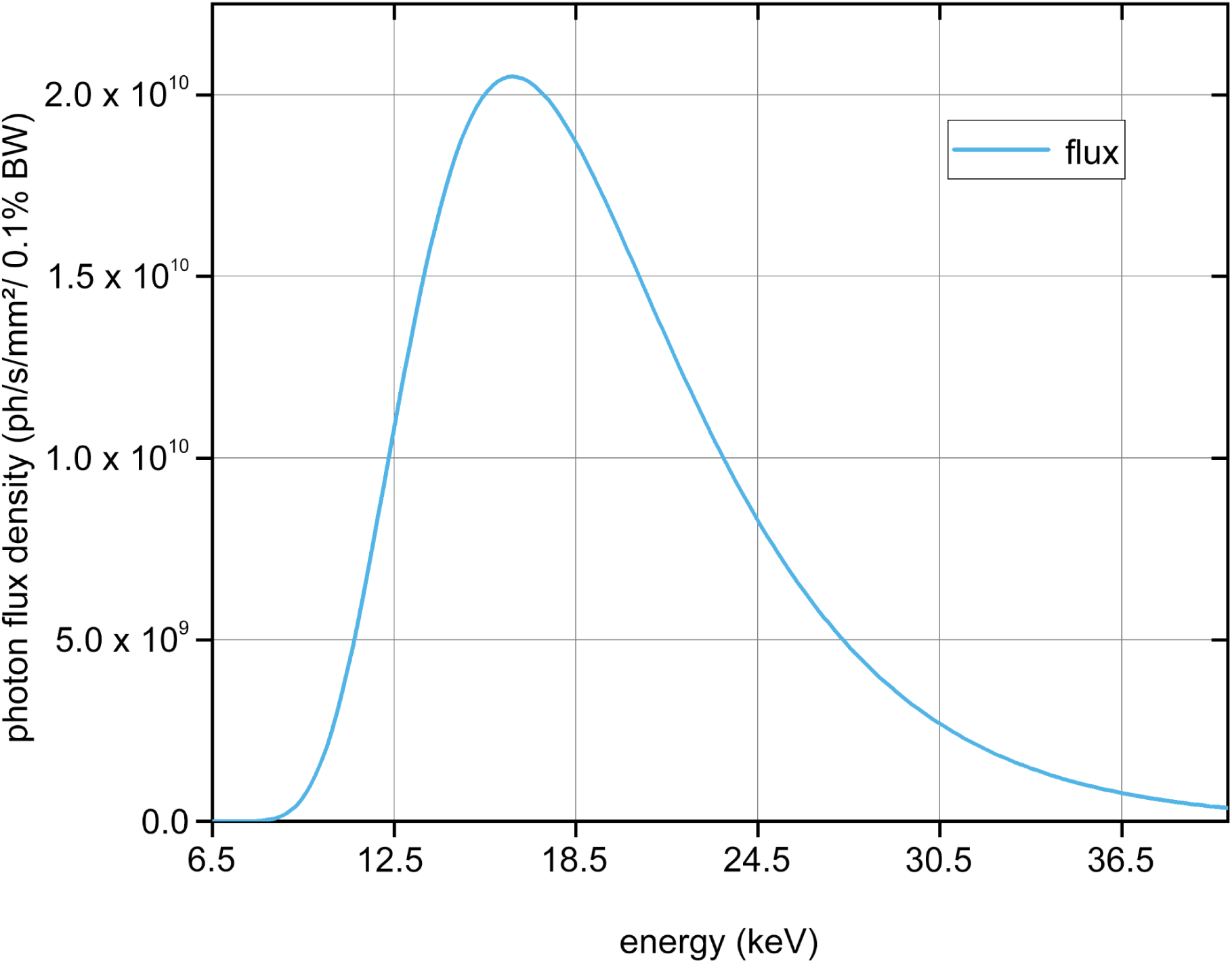
White beam spectrum with 10 mm PG filters. Peak at 16.5 keV, full width at half maximum of approximately 11 keV.

**Fig. S7.**
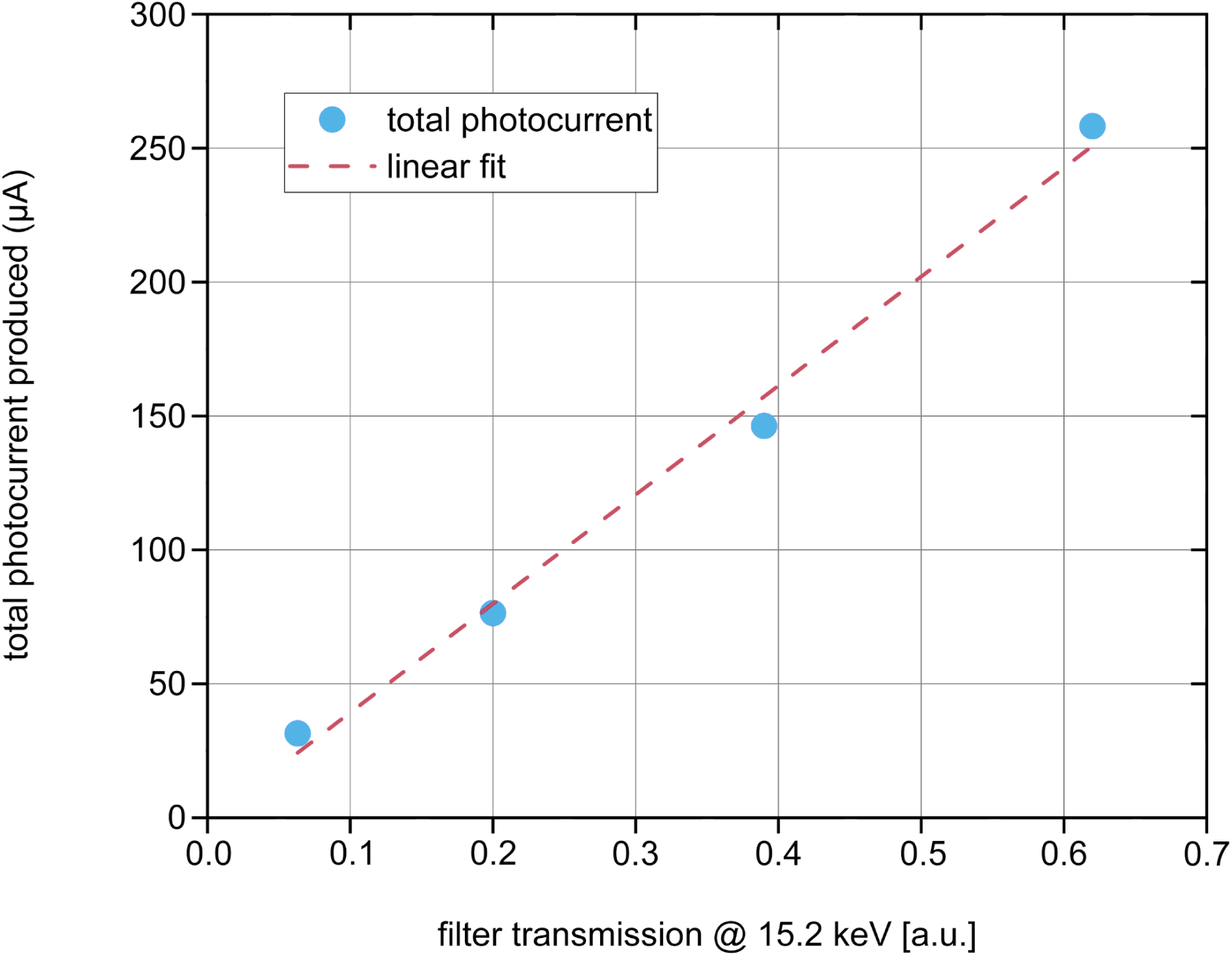
Total photocurrent generated in the pin diode by varying the impinging photon flux density.

**Fig. S8.**
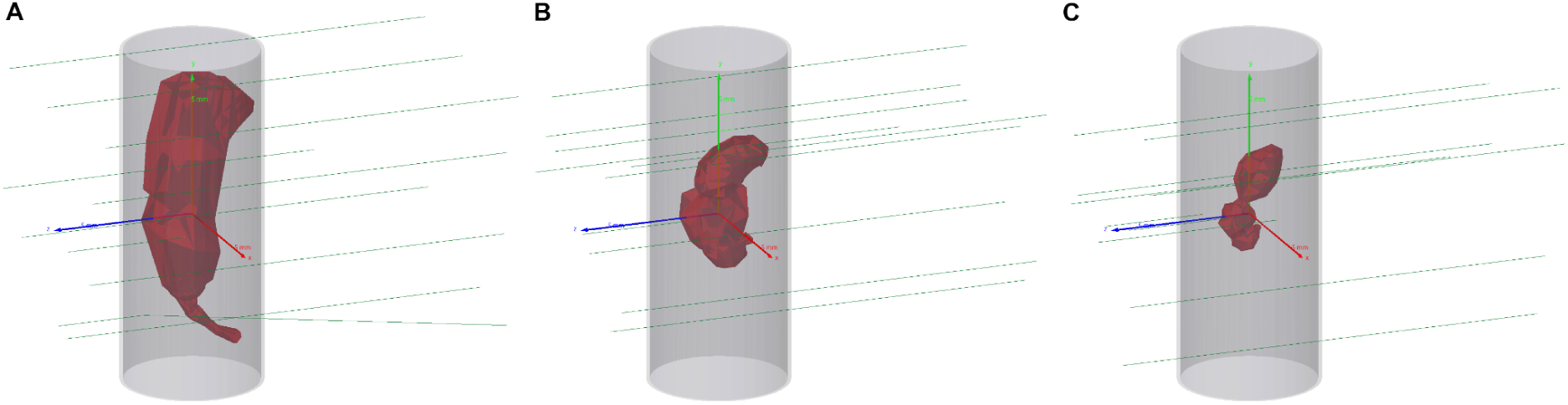
Simplified insect meshes and simulated particle tracks. a: *Sitophilus granarius*, b: *Drosophila suzukii*, c: *Leptopilina japonica*.

**Table S1.** Dose simulation results with a monochromatic 15.2 keV beam and filtered white beam at 125 mA ring current.

| Sample | Dose rate monochromatic (Gy/s) | Dose rate filtered white-beam (Gy/s) |
| --- | --- | --- |
| *Drosophila suzukii* | 189 ± 21 | 2742 ± 307 |
| *Leptopilina japonica* | 189 ± 21 | 2748 ± 307 |
| *Sitophilus granarius* | 191 ± 21 | 2770 ± 310 |

**Table S2.** Simulated dose in specimens exposed for various intervals at 125 mA ring current.

| Exposure time (seconds) | Dose monochromatic | Dose filtered white beam |
| --- | --- | --- |
| 2 ± 0.1 | 379 ± 46 Gy | 5.5 ± 0.7 kGy |
| 10 ± 0.1 | 1897 ± 213 Gy | 27.5 ± 0.3 kGy |
| 30 ± 0.1 | 5.7 ± 0.636 kGy | 82.6 ± 9.2 kGy |
| 60 ± 0.1 | 11.4 ± 1.3 kGy | 165.2 ± 18.5 kGy |
| 120 ± 0.1 | 22.8 ± 2.5 kGy | 330.4 ± 36.9 kGy |
| 300 ± 0.1 | 56.9 ± 6.4 kGy | 826.0 ± 92.3 kGy |
| 600 ± 0.1 | 113.8 ± 12.7 kGy | 1651.9 ± 184.7 kGy |
| 1200 ± 0.1 | 227.6 ± 25.4 kGy | 3303.8 ± 369.4 kGy |

**Table S3.** DNA concentration variance. Standard deviation values for the DNA concentration data of samples in EX1 and DNA concentration data of the negative control batch (no exposure specimens) in EX2.

|  | *S. granarius* | *D. suzukii* | *L. japonica* |
| --- | --- | --- | --- |
| HotSHOT (HT) | 2.333 | 1.049 | 0.592 |
| QuickExtract (QE) | 0.092 | 1.867 | 0.101 |
| Qiagen DNeasy (Qi) | 0.443 | 1.113 | 1.216 |
| Xiril (Xi) | 1.020 | 8.350 | 0.170 |
| Qiagen QIAamp (MU) | 3.964 | 4.327 | 4.399 |

**Table S4.** UCE loci. Proportion of UCE contigs that are >1 kb (contigs-L) in the full dataset of recovered loci.

|  | *S. granarius* | *D. suzukii* | *L. japonica* |
| --- | --- | --- | --- |
| Qiagen DNeasy (Qi) | 23.76% | 89.53% | 68.89% |
| Qiagen QIAamp (MU) | 52.12% | 89.68% | 81.09% |

**Dataset S1. Datasheet EX1.** Specimen datasheet for EX1. Experiment settings: Scanning mode and Extraction method, and Recorded data: Morphological evaluation and DNA concentration.

**Dataset S2. Datasheet EX2.** Specimen datasheet for EX2. Experiment settings: Exposure time, Ring current, Scanning mode and Extraction method, and Recorded data: DNA concentration, PCR & Sequencing success rates, Raw and Consensus sequence quality, Fragment length (PS and AS).

**Dataset S3. Fragment Analyser Data.** Raw electropherogram data for specimens in EX1 and EX2.

**Dataset S4. UCE data.** Raw UCE files and UCE contigs metrics: contigs-S, contigs-L, Total bp, mean length, 95 CI length, min length, max length, median length and Count.

**Dataset S5. Statistical analysis.** Output of statistical analysis performed on genetic data, faceted by taxon and integrated.

**Dataset S6. Statistical analysis: summary.** Output of statistical analysis performed on genetic data summarised for strongest factor, faceted by taxon, integrated and integrated: grouped factors.

**Movie S1. *Sitophilus granarius* before and after DNA extraction.** Virtual fly-through of tomographic datasets highlights the effects of different DNA extraction methods on internal anatomy.

**Movie S2. *Drosophila suzukii* before and after DNA extraction.** Virtual fly-through of tomographic datasets highlights the effects of different DNA extraction methods on internal anatomy.

**Movie S3. *Leptopilina japonica* before and after DNA extraction.** Virtual fly-through of tomographic datasets highlights the effects of different DNA extraction methods on internal anatomy.

## Notes

### Competing Interest Statement

The authors have declared no competing interest.

## References

1. Johnson, K. R., Owens, I. F., & Global Collection Group. (2023). A global approach for natural history museum collections. Science, 379(6638), 1192–1194.

2. Schowalter, T. D. (2022). Insect ecology: An ecosystem approach (5th ed.). Academic Press.

3. Scudder, G. G. E. (2017). The importance of insects. In R. G. Foottit & P. H. Adler (Eds.), Insect biodiversity.

4. Gorb, S. N., & Gorb, E. V. (2020). Insect-inspired architecture to build sustainable cities. Current Opinion in Insect Science, 40, 62–70.

5. Hallmann, C. A., Sorg, M., Jongejans, E., Siepel, H., Hofland, N., Schwan, H., … & De Kroon, H. (2017). More than 75 percent decline over 27 years in total flying insect biomass in protected areas. PloS One, 12(10), e0185809.

6. Chowdhury, S., Dubey, V. K., Choudhury, S., Das, A., Jeengar, D., Sujatha, B., … Kumar, V. (2023). Insects as bioindicator: A hidden gem for environmental monitoring. Frontiers in Environmental Science, 11, 1146052.

7. Irion, U., & Nüsslein-Volhard, C. (2022). Developmental genetics with model organisms. Proceedings of the National Academy of Sciences, 119(30), e2122148119.

8. Meier, R., Lawniczak, M. K., & Srivathsan, A. (2025). Illuminating entomological dark matter with DNA barcodes in an era of insect decline, deep learning, and genomics. Annual Review of Entomology, 70(1), 185–204.

9. Cabral-de-Mello, D. C., & Palacios-Gimenez, O. M. (2025). Repetitive DNAs: The ‘invisible’ regulators of insect adaptation and speciation. Current Opinion in Insect Science, 67, 101295.

10. Misof, B., Liu, S., Meusemann, K., Peters, R. S., Donath, A., Mayer, C., … & Zhou, X. (2014). Phylogenomics resolves the timing and pattern of insect evolution. Science, 346(6210), 763–767.

11. Li, F., Wang, X., & Zhou, X. (2025). The genomics revolution drives a new era in entomology. Annual Review of Entomology, 70(1), 379–400.

12. Watanabe, S., Masamura, N., Satoh, S. Y., & Hirao, T. (2024). Investigating the efficiency of DNA barcoding in insect classification: A review study. Entomology and Applied Science Letters, 1, 15–23.

13. Srivathsan, A., Ang, Y., Heraty, J. M., Hwang, W. S., Jusoh, W. F., Kutty, S. N., … & Meier, R. (2023). Convergence of dominance and neglect in flying insect diversity. Nature ecology & evolution, 7(7), 1012–1021.

14. Hotaling, S., Sproul, J. S., Heckenhauer, J., Powell, A., Larracuente, A. M., Pauls, S. U., … Frandsen, P. B. (2021). Long reads are revolutionizing 20 years of insect genome sequencing. Genome Biology and Evolution, 13(8), evab138.

15. Cameron, S. L. (2025). Insect mitochondrial genomics: A decade of progress. Annual Review of Entomology, 70(1), 83–101.

16. McCulloch, G. A., & Waters, J. M. (2023). Rapid adaptation in a fast-changing world: Emerging insights from insect genomics. Global Change Biology, 29(4), 943–954.

17. Short, A. E. Z., Dikow, T., & Moreau, C. S. (2018). Entomological collections in the age of big data. Annual Review of Entomology, 63, 513–530.

18. Lemmon, E. M., & Lemmon, A. R. (2013). High-throughput genomic data in systematics and phylogenetics. Annual Review of Ecology, Evolution, and Systematics, 44(1), 99–121.

19. Faircloth, B. C., McCormack, J. E., Crawford, N. G., Harvey, M. G., Brumfield, R. T., & Glenn, T. C. (2012). Ultraconserved elements anchor thousands of genetic markers spanning multiple evolutionary timescales. Systematic biology, 61(5), 717–726.

20. Blaimer, B. B., Lloyd, M. W., Guillory, W. X., & Brady, S. G. (2016). Sequence capture and phylogenetic utility of genomic ultraconserved elements obtained from pinned insect specimens. PLoS ONE, 11(8), e0161531.

21. Mera-Rodríguez, D., Fernández-Marín, H., & Rabeling, C. (2025). Phylogenomic approach to integrative taxonomy resolves a century-old taxonomic puzzle and the evolutionary history of the *Acromyrmex octospinosus* species complex. Systematic Entomology, 50(3), 469–494.

22. Prebus, M., & Rabeling, C. (2025). Phylogenomics resolve the systematics and biogeography of the ant tribe Myrmicini and tribal relationships within the hyperdiverse ant subfamily Myrmicinae. *Systematic Biology*, syaf022.

23. Lee, J.-Y. (2023). The principles and applications of high-throughput sequencing technologies. Development & Reproduction, 27(1), 9.

24. Blagoderov, V., Kitching, I. J., Livermore, L., Simonsen, T. J., & Smith, V. S. (2012). No specimen left behind: Industrial scale digitization of natural history collections. ZooKeys, 209, 133.

25. Ströbel, B., Schmelzle, S., Blüthgen, N., & Heethoff, M. (2018). An automated device for the digitization and 3D modelling of insects, combining extended-depth-of-field and all-side multi-view imaging. ZooKeys, (759), 1.

26. Poinapen, D., Konopka, J. K., Umoh, J. U., et al. (2017). Micro-CT imaging of live insects using carbon dioxide gas-induced hypoxia as anesthetic with minimal impact on certain subsequent life history traits. BMC Zoology, 2, 9.

27. Hall, M. J. R., & Martín-Vega, D. (2019). Visualization of insect metamorphosis. Philosophical Transactions of the Royal Society B, 374, 20190071.

28. Edel, C., Rühr, P. T., Frenzel, M., van de Kamp, T., Faragó, T., Hammel, J. U., Wilde, F., & Blanke, A. (2024). Bite force transmission and mandible shape in grasshoppers, crickets, and allies is not driven by dietary niches. Evolution, 78, 1958–1968.

29. Casadei-Ferreira, A., Procópio Camacho, G., van de Kamp, T., Lattke, J. E., Machado Feitosa, R., & Economo, E. P. (2025). Evolution and functional implications of stinger shape in ants. Evolution, 79(1), 80–99.

30. Soriano, C., Archer, M., Azar, D., Creaser, P., Delclòs, X., Godthelp, H., … & Tafforeau, P. (2010). Synchrotron X-ray imaging of inclusions in amber. Comptes Rendus Palevol, 9(6-7), 361–368.

31. Perreau, M., & Tafforeau, P. (2011). Virtual dissection using phase-contrast X-ray synchrotron microtomography: Reducing the gap between fossils and extant species. Systematic Entomology, 36, 573–580.

32. Rühr, P. T., van de Kamp, T., Faragó, T., Hammel, J., Wilde, F., Borisova, E., Edel, C., Frenzel, M., Baumbach, T., & Blanke, A. (2021). Juvenile ecology drives adult morphology in two insect orders. Proceedings of the Royal Society B, 288, 20210616.

33. Hoag, H. A., Raymond, M., Ulmer, J. M., Schwéger, S., van de Kamp, T., et al. (2025). The cranial gland system of *Nasonia* spp.: A link between chemical ecology, evo-devo, and descriptive taxonomy (Hymenoptera: Chalcidoidea). Journal of Insect Science, 25(2), ieaf034.

34. Püffel, F., Pouget, A., Liu, X., Zuber, M., van de Kamp, T., Roces, F., & Labonte, D. (2021). Morphological determinants of bite force capacity in insects: A biomechanical analysis of polymorphic leaf-cutter ants. Journal of the Royal Society Interface, 18, 20210424.

35. Ruan, Y., Konstantinov, A. S., Shi, G., Tao, Y., Li, Y., Johnson, A. J., … & Yang, X. (2020). The jumping mechanism of flea beetles (Coleoptera, Chrysomelidae, Alticini), its application to bionics and preliminary design for a robotic jumping leg. ZooKeys, 915, 87.

36. Betz, O., Wegst, U., Weide, D., Heethoff, M., Helfen, L., Lee, W. K., & Cloetens, P. (2007). Imaging applications of synchrotron X-ray phase-contrast microtomography in biological morphology and biomaterials science. I. General aspects of the technique and its advantages in the analysis of millimetre-sized arthropod structure. Journal of microscopy, 227(1), 51–71.

37. van de Kamp, T., Mikó, I., Staniczek, A. H., Eggs, B., Bajerlein, D., Faragó, T., … & Krogmann, L. (2022). Evolution of flexible biting in hyperdiverse parasitoid wasps. Proceedings of the Royal Society B: Biological Sciences, 289(1967).

38. Vommaro, M. L., Donato, S., Caputo, S., Agostino, R. G., Montali, A., Tettamanti, G., & Giglio, A. (2024). Anatomical changes of Tenebrio molitor and Tribolium castaneum during complete metamorphosis. Cell and Tissue Research, 396(1), 19–40.

39. van de Kamp, T., Schwermann, A. H., dos Santos Rolo, T., Lösel, P. D., Engler, T., Etter, W., … & Krogmann, L. (2018). Parasitoid biology preserved in mineralized fossils. Nature Communications, 9(1), 3325.

40. Katzke, J., Hita Garcia, F., Lösel, P. D., Azuma, F., Faragó, T., Aibekova, L., … & van de Kamp, T. (2026). High-throughput phenomics of global ant biodiversity. Nature Methods.

41. Jonsson, T. (2023). Micro-CT and deep learning: Modern techniques and applications in insect morphology and neuroscience. Frontiers in Insect Science, 3, 1016277.

42. Roots, R., & Okada, S. (1975). Estimation of life times and diffusion distances of radicals involved in X-ray-induced DNA strand breaks or killing of mammalian cells. Radiation Research, 64, 306–320.

43. Sutherland, B. M., Bennett, P. V., Sutherland, J. C., & Laval, J. (2002). Clustered DNA damages induced by x rays in human cells. Radiation research, 157(6), 611–616.

44. Parplys, A. C., Petermann, E., Petersen, C., Dikomey, E., & Borgmann, K. (2012). DNA damage by X-rays and their impact on replication processes. Radiotherapy and oncology, 102(3), 466–471.

45. Hiszczynska-Sawicka, E., Richards, N. K., Gibson, J. L., van Koten, C., Gunawardana, D., & Armstrong, K. F. (2025). Defining the parameters of comet assay for reliably detecting DNA damage as a marker of X-ray based irradiation in insects treated with non-lethal doses for export fruit sanitation. Pest Management Science, 81(10), 7232–7243.

46. Truett, G. E., Heeger, P., Mynatt, R. L., Truett, A. A., Walker, J. A., & Warman, M. L. (2000). Preparation of PCR-quality mouse genomic DNA with hot sodium hydroxide and tris (HotSHOT). BioTechniques, 29(1), 52–54.

47. Feng, V., Høegh-Guldberg, C., Meier, R., & Buček, A. (2026). Balancing Barcoding and Genomics: gDNA Quality in Insect Vouchers After HotSHOT DNA Extraction. Molecular Ecology Resources, 26(2), e70103.

48. Mouzaki, D. G., & Margaritis, L. H. (1994). The eggshell of the almond wasp *Eurytoma amygdali* (Hymenoptera, Eurytomidae): Morphogenesis and fine structure of the eggshell layers. Tissue and Cell, 26(4), 559–568.

49. Crowther, R. A., DeRosier, D. J., & Klug, A. (1970). The reconstruction of a three-dimensional structure from projections and its application to electron microscopy. Proceedings of the Royal Society of London. A. Mathematical and Physical Sciences, 317(1530), 319–340.

50. Cecilia, A., Simon, R., Hamann, E., Zuber, M., Faragó, T., Haenschke, D., … & Baumbach, T. (2025). The IMAGE beamline at the KIT Light Source. Synchrotron Radiation, 32(4).

51. Cecilia, A., Rack, A., Douissard, P. A., Martin, T., dos Santos Rolo, T., Vagovič, P., … & Baumbach, T. (2011). LPE grown LSO: Tb scintillator films for high-resolution X-ray imaging applications at synchrotron light sources. Nuclear instruments and methods in physics research section A: Accelerators, spectrometers, detectors and associated equipment, 648, S321–S323.

52. Douissard, P. A., Cecilia, A., Rochet, X., Chapel, X., Martin, T., van de Kamp, T., … & Rack, A. (2012). A versatile indirect detector design for hard X-ray microimaging. Journal of Instrumentation, 7(09), P09016–P09016.

53. Paganin, D., Mayo, S. C., Gureyev, T. E., Miller, P. R., & Wilkins, S. W. (2002). Simultaneous phase and amplitude extraction from a single defocused image of a homogeneous object. Journal of Microscopy, 206, 33–40.

54. Vogelgesang, M., Chilingaryan, S., dos Santos Rolo, T., & Kopmann, A. (2012). High performance computing techniques for tomographic reconstruction. In 2012 IEEE 14th International Conference on High Performance Computing and Communication & 2012 IEEE 9th International Conference on Embedded Software and Systems (pp. 824–829). IEEE.

55. Faragó, T., Gasilov, S., Emslie, I., Zuber, M., Helfen, L., Vogelgesang, M., & Baumbach, T. (2022). Tofu: a fast, versatile and user-friendly image processing toolkit for computed tomography. Synchrotron Radiation, 29(3), 916–927.

56. Limaye, A. (2012). Drishti: A volume exploration and presentation tool. Proceedings of SPIE, 8506, 85060X.

57. Lösel, P. D., van de Kamp, T., Jayme, A., Ershov, A., Faragó, T., Pichler, O., … & Heuveline, V. (2020). Introducing Biomedisa as an open-source online platform for biomedical image segmentation. Nature communications, 11(1), 5577.

58. Ban, S., Hirayama, H., Namito, Y., Tanaka, S., Nakashima, H., Nakane, Y., & Nariyama, N. (1994). Calibration of silicon PIN photodiode for measuring intensity of 7–40 keV photons. Journal of Nuclear Science and Technology, 31(2), 163–168.

59. Agostinelli, S., Allison, J., Amako, K., Apostolakis, J., Araujo, H., Arce, P., … & Zschiesche, D. (2003). Geant4—a simulation toolkit. Nuclear Instruments and Methods in Physics Research Section A: Accelerators, Spectrometers, Detectors and Associated Equipment, 506, 250–303.

60. Allison, J., Amako, K., Apostolakis, J. E. A., Araujo, H. A. A. H., Dubois, P. A., Asai, M. A. A. M., … & Yoshida, H. A. Y. H. (2006). Geant4 developments and applications. IEEE Transactions on nuclear science, 53(1), 270–278.

61. Allison, J., Amako, K., Apostolakis, J., Arce, P., Asai, M., Aso, T., … & Yoshida, H. (2016). Recent developments in Geant4. Nuclear instruments and methods in physics research section A: Accelerators, Spectrometers, Detectors and Associated Equipment, 835, 186–225.

62. Fagan, W. F., Siemann, E., Mitter, C., Denno, R. F., Huberty, A. F., Woods, H. A., & Elser, J. J. (2002). Nitrogen in insects: implications for trophic complexity and species diversification. The American Naturalist, 160(6), 784–802.

63. Back, J. A., & King, R. S. (2013). Sex and size matter: Ontogenetic patterns of nutrient content of aquatic insects. Freshwater Science, 32, 837–848.

64. Hackman, R. H. (1974). Chemistry of the arthropod cuticle. In M. Rockstein (Ed.), The physiology of Insecta (2nd ed., pp. 215–270). Academic Press.

65. Srivathsan, A., Lee, L., Katoh, K., Hartop, E., Kutty, S. N., Wong, J., … Meier, R. (2021). ONTbarcoder and MinION barcodes aid biodiversity discovery and identification by everyone, for everyone. BMC Biology, 19(1), 217.

66. Vasilița, C., Bremer, J., Popovici, O. A., Krogmann, L., & Talamas, E. (in press). From shadows to clarity: Resolving the taxonomy of *Gryon* (Hymenoptera: Scelionidae) by integrating modern and historic data. Proceedings of the Entomological Society of Washington.

67. Haas, M., Baur, H., Schweizer, T., Monje, J. C., Moser, M., Bigalk, S., & Krogmann, L. (2021). Tiny wasps, huge diversity–A review of German Pteromalidae with new generic and species records (Hymenoptera: Chalcidoidea). Biodiversity Data Journal, 9, e77092.

68. Ivanova, N. V., Dewaard, J. R., & Hebert, P. D. (2006). An inexpensive, automation-friendly protocol for recovering high-quality DNA. Molecular Ecology Notes, 6(4), 998–1002.

69. Cruaud, A., Nidelet, S., Arnal, P., Weber, A., Fusu, L., Gumovsky, A., … & Rasplus, J. Y. (2019). Optimized DNA extraction and library preparation for minute arthropods: Application to target enrichment in chalcid wasps used for biocontrol. Molecular Ecology Resources, 19(3), 702–710.

70. Lutgen, D., & Burri, R. (2022). DNA extraction protocol for historical toe pad samples from birds. Protocols.io. 10.17504/protocols.io.bm4mk8u6

71. Folmer, O., M. Black, W. Hoeh, R. Lutz and R. Vrijenhoek. 1994. DNA primers for amplification of mitochondrial Cytochrome C oxidase subunit I from diverse metazoan invertebrates. Molecular Marine Biology and Biotechnology 3: 294–299.

72. Ratnasingham, S., & Hebert, P. D. (2007). BOLD: The Barcode of Life Data System. Molecular Ecology Notes, 7, 355–364.

73. Fisher, S., Barry, A., Abreu, J., Minie, B., Nolan, J., Delorey, T. M., … & Nusbaum, C. (2011). A scalable, fully automated process for construction of sequence-ready human exome targeted capture libraries. Genome biology, 12(1), R1.

74. Gnirke, A., Melnikov, A., Maguire, J., Rogov, P., LeProust, E. M., Brockman, W., … & Nusbaum, C. (2009). Solution hybrid selection with ultra-long oligonucleotides for massively parallel targeted sequencing. Nature biotechnology, 27(2), 182–189.

75. Faircloth, B. C., Branstetter, M. G., White, N. D., & Brady, S. G. (2015). Target enrichment of ultraconserved elements from arthropods provides a genomic perspective on relationships among Hymenoptera. Molecular ecology resources, 15(3), 489–501.

76. Bankevich, A., Nurk, S., Antipov, D., Gurevich, A. A., Dvorkin, M., Kulikov, A. S., … & Pevzner, P. A. (2012). SPAdes: a new genome assembly algorithm and its applications to single-cell sequencing. Journal of computational biology, 19(5), 455–477.

77. Box, G. E., & Cox, D. R. (1964). An analysis of transformations. Journal of the Royal Statistical Society Series B: Statistical Methodology, 26(2), 211–243.

78. Shapiro, S. S., & Wilk, M. B. (1965). An analysis of variance test for normality (complete samples). Biometrika, 52(3-4), 591–611.

79. Levene, H. (1960). Robust tests for equality of variances. Contributions to probability and statistics, 278–292.

80. Kruskal, W. H., & Wallis, W. A. (1952). Use of ranks in one-criterion variance analysis. Journal of the American Statistical Association, 47(260), 583–621.

81. Tukey, J. W. (1949). Comparing individual means in the analysis of variance. Biometrics, 99–114.

82. Dunn, O. J. (1961). Multiple comparisons among means. Journal of the American statistical association, 56(293), 52–64.

83. Pedregosa, F., Varoquaux, G., Gramfort, A., Michel, V., Thirion, B., Grisel, O., … & Duchesnay, É. (2011). Scikit-learn: Machine learning in Python. The Journal of Machine Learning Research, 12, 2825–2830.

84. Virtanen, P., Gommers, R., Oliphant, T. E., Haberland, M., Reddy, T., Cournapeau, D., … & Van Mulbregt, P. (2020). SciPy 1.0: fundamental algorithms for scientific computing in Python. Nature Methods, 17(3), 261–272.

85. Aton, M., McDonald, D., Cañardo Alastuey, J., Azom, R., Batra, P., Bezshapkin, V., … & Zhu, Q. (2025). Scikit-bio: a fundamental Python library for biological omic data analysis. Nature Methods, 1–3.

86. Hunter, J. D. (2007). Matplotlib: A 2D graphics environment. Computing in science & engineering, 9(3), 90–95.

87. Waskom, M. L. (2021). Seaborn: statistical data visualization. Journal of open source software, 6(60), 3021.

